# Massively parallel disruption of enhancers active during human corticogenesis

**DOI:** 10.1101/852673

**Authors:** Evan Geller, Jake Gockley, Deena Emera, Severin Uebbing, Justin Cotney, James P. Noonan

## Abstract

Changes in gene regulation have been linked to the expansion of the human cerebral cortex and to neurodevelopmental disorders. However, the biological effects of genetic variation within developmental regulatory elements on human corticogenesis are not well understood. We used sgRNA-Cas9 genetic screens in human neural stem cells (hNSCs) to disrupt 10,674 expressed genes and 2,227 enhancers active in the developing human cortex and determine the resulting effects on cellular proliferation. Gene disruptions affecting proliferation were enriched for genes associated with risk for human neurodevelopmental phenotypes including primary microcephaly and autism spectrum disorder. Although disruptions in enhancers had overall weaker effects on proliferation than gene disruptions, we identified enhancer disruptions that severely perturbed hNSC self-renewal. Disruptions in Human Accelerated Regions and Human Gain Enhancers, regulatory elements implicated in the evolution of the human brain, also perturbed hNSC proliferation, establishing a role for these elements in human neurodevelopment. Integrating proliferation phenotypes with chromatin interaction maps revealed regulatory relationships between enhancers and target genes that contribute to neurogenesis and potentially to human cortical evolution.

## Main Text

The development of the human cerebral cortex depends on the precise spatial, temporal and quantitative control of gene expression by transcriptional enhancers (*1*). Genetic variants with the potential to alter gene expression in the developing brain have been implicated both in neurodevelopmental disorders and in the expansion and elaboration of the cortex during human evolution (*2–10*). Despite the growing evidence that regulatory variation influences human brain phenotypes, the biological effects of genetic changes that occur within enhancers active during human corticogenesis have not been systemically studied.

Here we employ a massively parallel sgRNA-Cas9 genetic screen in H9-derived human neural stem cells (hNSCs) to disrupt 2,227 enhancers known to be active during human corticogenesis and identify enhancers required for normal hNSC proliferation (Fig. 1A) (*11–14*). We chose hNSCs as a model of corticogenesis because of the fundamental role of the neural stem cell niche in neurogenesis and the specification of cortical size (*15, 16*). The hNSCs we used express multiple neural stem cell markers, including *NES, SOX2*, *PAX6*, and *HES1*, and are multipotent, capable of differentiating into neurons, oligodendrocytes and astrocytes (supplementary text) (*17, 18*). Regulatory variants that alter neural stem cell proliferation and self-renewal could result in changes to the number, type and proportion of cortical neurons generated during cortical development, and these changes may contribute to disorders of brain development and function (*17*). In addition, the human cortex exhibits a greatly expanded number of progenitor cells during development compared to other primates, suggesting that modification of hNSC proliferation contributed to the increase in cortical size during human evolution (*16, 17*).

**Fig. 1.**
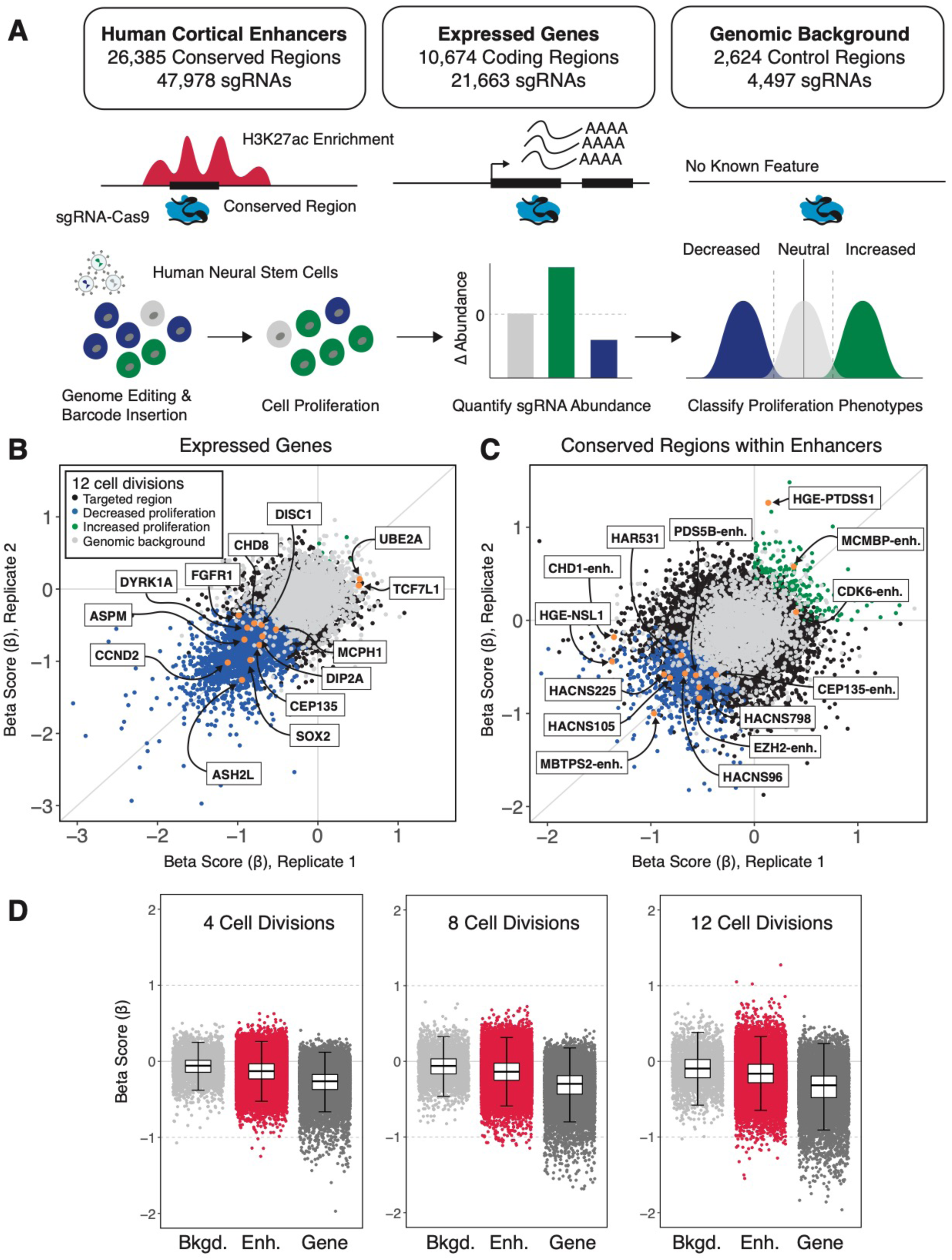
Identifying genes and enhancers that alter human neural stem cell (hNSC) proliferation following genetic disruption. (**A**) (*Top*) Summary of conserved regions within enhancers, genes, and background control regions targeted in the sgRNA-Cas9 genetic screen. Putative transcription factor binding sites within enhancers were identified based on the placental mammal PhastCons element annotation (GRCh37/hg19) (*24*). Each PhastCons element was divided into ∼30 base-pair conserved regions and targeted for genetic disruption by sgRNAs. Genes were targeted based on evidence of expression in hNSCs. Genomic background controls are non-coding regions with no detectable evidence of regulatory activity based on epigenetic signatures (see supplement). (*Bottom*) Schematic illustrating transduction of sgRNA-Cas9 lentiviral libraries into hNSCs at a low multiplicity of infection (MOI ∼0.3) to ensure that cells were infected by a single virion and that ∼1,000 cells were infected per sgRNA. Changes in the abundances of each sgRNA over time and classification of proliferation phenotypes was carried out as described in the main text. **(B, C)** Scatterplots illustrating Beta scores (β) across two biological replicates resulting from genetic disruption of genes (**B**) and enhancers (**C**), measured at 12 cell divisions following transduction. The effect of each gene or enhancer disruption is indicated in the legend shown in **B**. Gene and enhancer disruptions affecting proliferation in regions with known biological roles in neural stem cells are identified by name (*orange*). Examples of genes were selected based on neural stem cell maintenance or association with neurodevelopmental disorders. Enhancer examples were selected based on evidence of a regulatory interaction with genes with roles in chromatin modification and association with neurodevelopmental disorders or their identification as a HAR or HGE. (**D**) Distribution of proliferation phenotypes resulting from disruption of genomic background control regions (*light grey*), enhancers (*red*) and genes (*dark grey*) at 4, 8 and 12 cell divisions. Box plots illustrate the lower (25th percentile), median (50th percentile) and upper (75th percentile) of Beta scores (β) for each category. Error bars indicate an estimated 95% confidence interval.

To disrupt regions likely to encode putative transcription factor binding sites within enhancers, we targeted 26,385 conserved regions (47,978 total sgRNAs) across the 2,227 enhancers included in our screen. We selected these enhancers based on two criteria. First, the enhancers are marked by H3K27ac, a histone modification associated with enhancer activity, both in hNSCs and in human cortex during developmental periods that include the expansion of proliferative zones and the onset of neurogenesis (*2, 18*). Second, we filtered these enhancers for evidence of strong H3K27ac marking in the developing cortex compared to other human tissues, in order to enrich for enhancers with cortical-specific functions (fig. S1; supplementary text).

We also included in our screen two classes of enhancers implicated in human brain evolution: 93 Human Accelerated Regions (HARs) and 129 Human Gain Enhancers (HGEs). HARs harbor an excess of human-specific substitutions and exhibit human-specific activities during development (*4–7, 19*). HGEs have increased levels of enhancer-associated chromatin marks in the developing human cortex relative to other species (*2, 20*). Chromosome conformation studies in the developing cortex suggest that both HARs and HGEs interact with genes involved in neurogenesis, axon guidance and synaptic transmission (*21–23*). However, a functional role for HARs and HGEs in regulating human neurogenesis has not been established.

To directly compare the effects of enhancer and gene disruptions, we also targeted 10,674 protein-coding genes (21,663 sgRNAs) actively transcribed in hNSCs (Fig. 1A). We included 2,624 genomic background regions (4,497 sgRNAs) and 500 non-targeting sgRNA controls to respectively account for non-specific effects of inducing small genetic lesions that require repair and background effects of lentiviral transduction (supplementary text). We defined genomic background regions as non-coding sequences that exhibited no evidence of function based on epigenetic marking in human tissues and cells (supplementary text). In total, this yielded a library of 74,138 sgRNAs that we transduced into hNSCs across 8 sub-libraries (fig. S2). The abundance of each integrated sgRNA was determined using PCR followed by high-throughput sequencing, initially after lentiviral transduction, and then subsequently at 4, 8 and 12 cell divisions (Fig. 1A; supplementary text). Modeling the change in abundance of each sgRNA across the time series provided a quantitative basis for measuring effects on cellular proliferation (*25*). We hypothesized that alterations in hNSC proliferation would encompass diverse cellular changes, including disrupted cell cycle regulation, differentiation of hNSCs into derived cell types, and reduced cell survival.

We first quantified the biological effects of targeted disruption on hNSC proliferation, the Beta score (β), for each gene, conserved region, or genomic background control relative to a set of non-targeting controls (Fig. 1B-D; supplementary text). These biological effects were determined using reproducible sgRNA read abundances collected across both replicates and multiple time points (Pearson correlation > 0.9) (fig. S3-5; table S1) and demonstrate high levels of on-target specificity (fig. S6; supplementary text). We then performed linear discriminant analysis (LDA) to partition gene, enhancer, and background control disruptions into proliferation-decreasing, proliferation-increasing, or neutral classes (fig. S7; supplementary text). We performed this analysis on a training set including known proliferation-decreasing genes, background controls, and the top proliferation-increasing effects across time points (empirical FDR < 0.05) (*26*). The trained classifier was then applied to the full dataset and filtered for consistency in classification and the direction of the effect across timepoints to identify genetic disruptions resulting in proliferation phenotypes (Fig. 1A; supplementary text).

We identified 2,263 genes (22.9% of all targeted genes) that alter hNSC proliferation at 12 cell divisions (Fig.1B-D; Table 1). Of these, nearly all gene disruption phenotypes showed decreased proliferation, while only 8 gene disruptions increased proliferation (Fig. 1B; Table 1). Many gene disruptions that altered proliferation have known roles in neural stem cell biology relating to the balance between self-renewal and neuronal differentiation (e.g., *CCND2*, *SOX2*), or response to growth factor signaling (e.g., *FGFR1*, *TCF7L1*) (Fig. 1B) (*27–30*). Disruption of genes associated with microcephaly (e.g., *ASPM*, *CEP135*, *MCPH1)* decreased hNSC proliferation, consistent with their known roles in human cortical development (Fig. 1B) (*31*). Disruptions of genes associated with risk for other neurodevelopmental disorders, notably autism spectrum disorder (e.g., *DYRK1A*, *DIP2A*, *CHD8*) and X-linked intellectual disability (e.g., *UBE2A*), also resulted in significant alterations to hNSC proliferation (*31–33*).

**Table 1.**
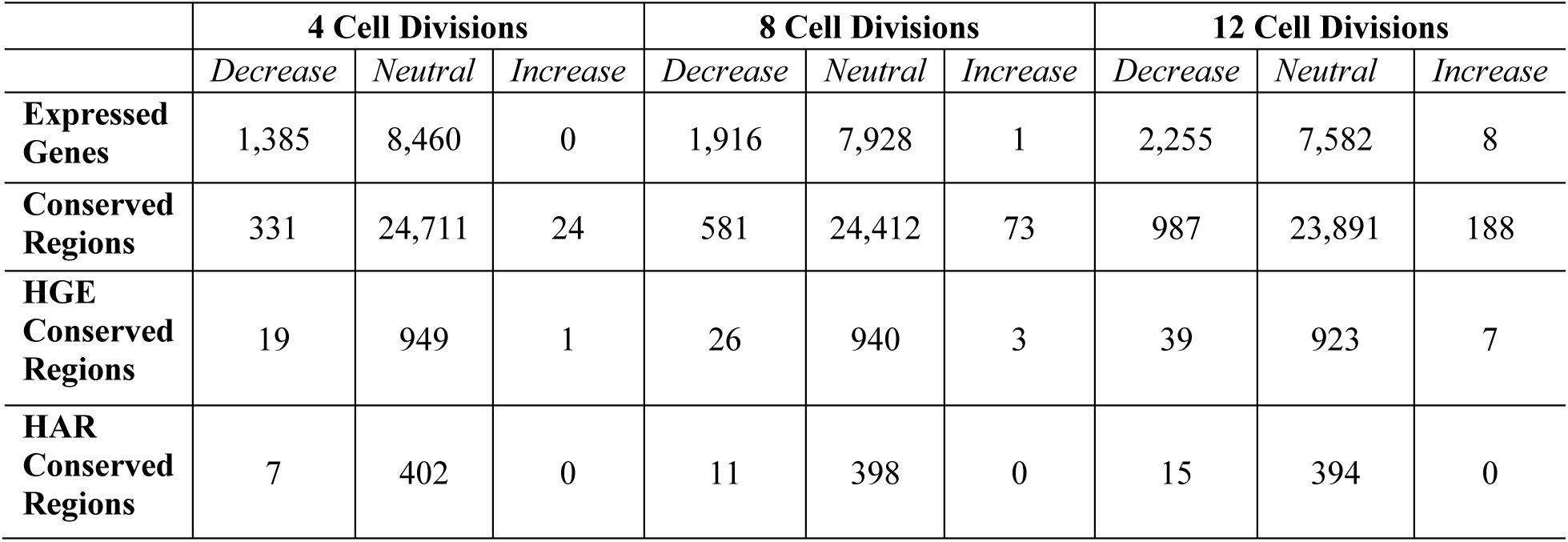
Observed proliferation phenotypes for gene and enhancer disruptions in hNSCs.

|  | 4 Cell Divisions |  |  | 8 Cell Divisions |  |  | 12 Cell Divisions |  |  |
| --- | --- | --- | --- | --- | --- | --- | --- | --- | --- |
|  | *Decrease* | *Neutral* | *Increase* | *Decrease* | *Neutral* | *Increase* | *Decrease* | *Neutral* | *Increase* |
| **Expressed Genes** | 1,385 | 8,460 | 0 | 1,916 | 7,928 | 1 | 2,255 | 7,582 | 8 |
| **Conserved Regions** | 331 | 24,711 | 24 | 581 | 24,412 | 73 | 987 | 23,891 | 188 |
| **HGE Conserved Regions** | 19 | 949 | 1 | 26 | 940 | 3 | 39 | 923 | 7 |
| **HAR Conserved Regions** | 7 | 402 | 0 | 11 | 398 | 0 | 15 | 394 | 0 |

We found 1,175 conserved regions (4.9% of all conserved regions) across 708 cortical enhancers (31.7% of all targeted enhancers) that alter hNSC proliferation by 12 cell divisions. (Fig. 1B-D; Table 1). In contrast to genes, a greater proportion (16%) of disruptions in enhancers increased proliferation (188 proliferation-increasing versus 987 proliferation-decreasing; Table 1). We also discovered 46 conserved regions within HGEs and 15 within HARs that alter proliferation when disrupted (Fig. 1C, Table 1), supporting that HARs and HGEs contribute to human neurodevelopment. Disruption of HGEs affecting proliferation were identified as proximal to genes with molecular functions in chromosome segregation *(e.g., NSL1)* or associated with intellectual disability (e.g., *PTDSS1)* (*34, 35*). Proliferation phenotypes identified within three HARs altered conserved regions that contained human-specific substitutions; these HARs were located in introns of genes with known functions in brain development *(e.g., NPAS3*) and cognitive function (e.g., *USH2A*) (Fig. 1C) (*36, 37*).

Globally, disruptions within enhancers had quantitatively weaker effects on proliferation than gene disruptions (Fig. 1D) (Wilcoxon rank sum test, *P* < 2.2×10^-16^). Although many enhancer disruptions resulted in biological effects of comparable magnitude to gene disruptions, we observed differences in the onset of their biological effects (Fig. 1D). The majority of gene disruption phenotypes (61% of total gene phenotypes) were detected by 4 cell divisions. In contrast, fewer enhancer disruption phenotypes (30% of all conserved region phenotypes) were detected at this early timepoint. The total number of phenotypes within enhancers approximately doubled at 8 cell divisions and doubled again at 12 cell divisions (Table 1). The later onset of enhancer phenotypes may be due to delayed effects on target gene expression, slow turnover of transcripts and protein products, or buffering of effects on gene expression due to partial redundancy among multiple enhancers regulating a single target gene.

Measuring the effect of gene and enhancer disruptions across multiple timepoints allowed us to distinguish the overall effect on proliferation from temporal changes across cell divisions. We used principal component analysis (PCA) to extract these latent factors and found that the first principal component (PC1 = 94.3% of total variance) is correlated with the severity of the effect on cellular proliferation (Fig. 2A; fig. S8). This analysis enabled us to assign a proliferation score to each disruption, which we could then use to rank disruptions based on their cumulative effect on cellular proliferation across multiple timepoints. The second component (PC2 = 3.9% of total variance) correlated with effect changes across time points. Examples include the continued increase in proliferation resulting from genetic disruption of the X-linked intellectual disability gene *UBE2A* (Fig. 2A) (*33*) and the decrease in proliferation due to disruption of *KIF20B* (Fig. 2A), a gene implicated in microcephaly (*31*). These findings support that both proliferation-increasing and proliferation-decreasing phenotypes revealed by massively parallel screening in hNSCs provide insight into human neurodevelopmental disorders. Together, these latent factors explain nearly all of the variability in the biological effects of perturbation to genes and enhancers (>98% of total variance) and we utilized these factors to dissect the functional characteristics of hNSC proliferation (Fig. 2A; table S2).

**Fig. 2.**
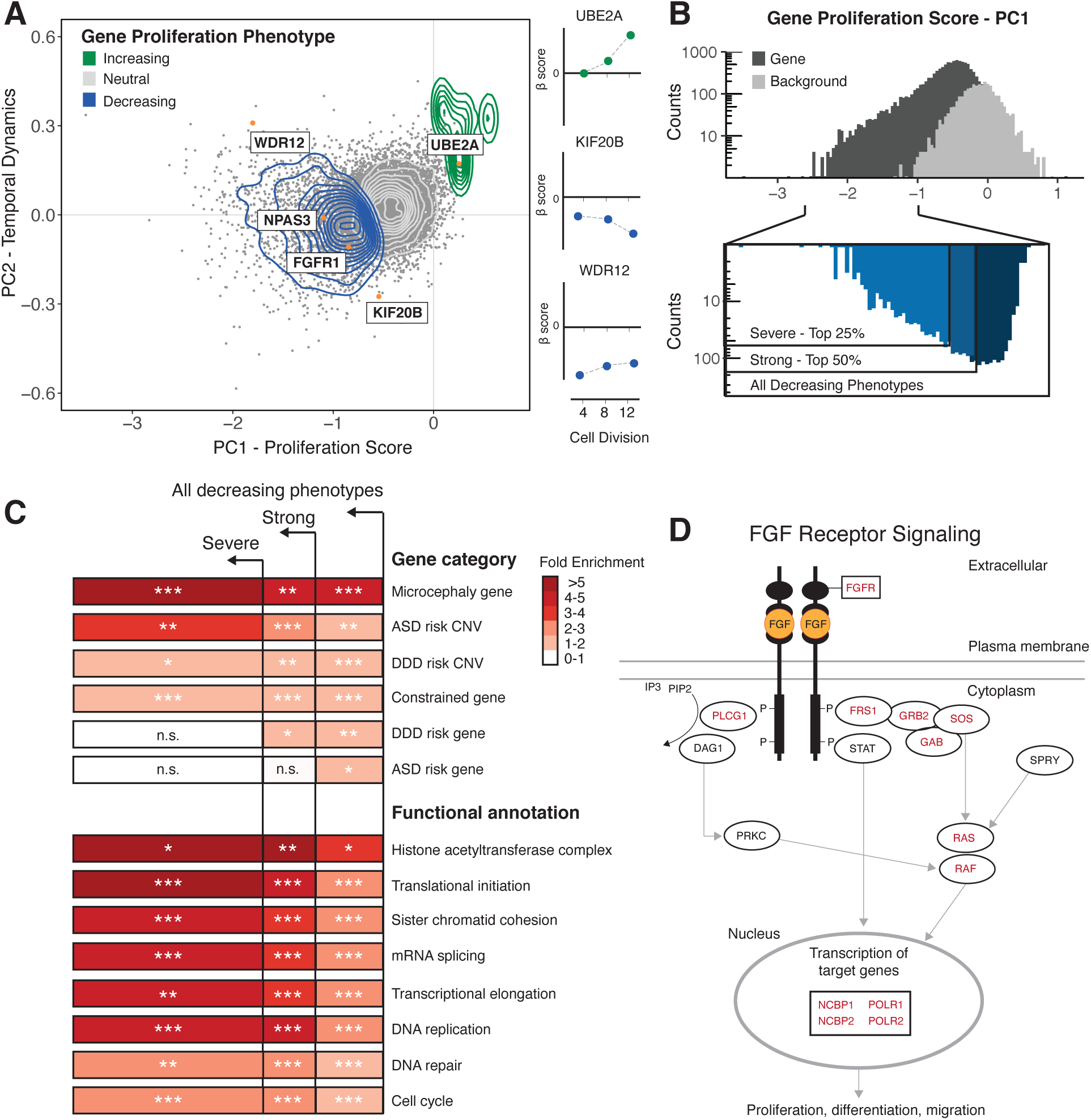
Characterization of gene proliferation phenotypes. (**A**) (*Left*) Principal component analysis (PCA) of all gene disruptions, with a 2-D density overlay illustrating the distribution of proliferation-decreasing (*blue*), neutral (*grey*), and proliferation-increasing (*green*) phenotypes. Individual gene disruptions (*orange*) with biological roles in the maintenance of neural stem cells or human neurodevelopment (*31, 33, 41*). (*Right)* Temporal dynamics captured by principal component analysis shown at 4, 8, and 12 cell divisions. (**B**) (*Top*) Histogram of proliferation scores obtained from PCA for gene disruptions (*black*) and genomic background controls (*grey*). (*Bottom*) Partitioning of all gene disruptions that decrease proliferation into severe (top 25%) and strong (top 50%) categories. (**C**) Fold-enrichment of gene ontology biological processes and risk gene sets within the partitions shown in (**B)**. n.s. = not significant, * = hypergeometric BH-corrected p-value < 0.05; ** = hypergeometric BH-corrected p-value < 0.005; *** = hypergeometric BH-corrected p-value < 0.0005. (**D**) FGF signaling pathway depicted by Reactome (*42*) and the literature (*43*). Genes disrupted within this pathway are shown in black, and a significant subset result in proliferation phenotypes (in red).

We first examined whether gene disruptions converge on specific biological pathways and known human disease phenotypes. We hypothesized that gene disruptions with stronger effects might be functionally distinct from disruptions with weaker effects. We therefore grouped the proliferation phenotypes into categories based on their proliferation scores and performed overrepresentation testing of proliferation phenotypes across developmental disorder risk loci, Gene Ontology biological processes, and biological signaling pathways (Fig. 2B-D; table S3-S5). We found that genes associated with neurodevelopmental disorders were significantly overrepresented among gene disruptions that altered proliferation. Genes associated with microcephaly were enriched in all three categories and were most strongly enriched in the most severe set (hypergeometric test, *P* = 3.4 × 10^-4^) (Fig. 2C). This is consistent with known disease processes that impair cell division in the developing human cortex (*31*). Genes located within large copy-number variants (CNVs) associated with autism spectrum disorders (ASD) (*32*) also showed strong enrichment in severe phenotypes (hypergeometric test, *P* = 1.1 × 10^-3^), providing the means to identify new potential candidate genes in these regions (table S3). CNVs associated with developmental disorders (hypergeometric test, *P* = 1.6 × 10^-3^) (*38*), as well as constrained genes intolerant to loss-of-function mutations (hypergeometric test, *P* = 5.1 × 10^-17^) exhibited moderate enrichment across all phenotypes (Fig. 2C) (*39*). Finally, many risk genes associated with developmental disorders and ASD have been identified based on a significant excess of gene disrupting loss-of-function mutations in affected individuals (*32, 40*). These genes are also significantly enriched among proliferation-altering gene disruptions (hypergeometric test, *P* = 3.2 × 10^-6^ for genes associated with developmental disorders and hypergeometric test, *P* = 9.8 × 10^-3^ for genes associated with ASD) (Fig. 2C), suggesting that loss-of-function mutations in these genes may contribute to developmental disorders in part by perturbing hNSC proliferation.

We then used Gene Ontology analysis to identify biological processes enriched among proliferation-altering gene disruptions (table S4). In severe proliferation phenotypes, we observed the strongest fold-enrichment for gene functions related to histone acetyltransferase activity (modified Exact test, BH-corrected *P* = 2.1 × 10^-3^) and translational initiation (modified Exact test, BH-corrected *P* = 1.4 × 10^-12^) (Fig. 2C). Additional processes exhibiting elevated fold-enrichment in severe phenotypes include sister chromatid cohesion (modified Exact test, BH-corrected *P* = 1.7 × 10^-6^), mRNA splicing (modified Exact test, BH-corrected *P* = 4.3 × 10^-12^), transcriptional elongation (modified Exact test, BH-corrected *P* = 2.4 × 10^-5^), and DNA replication (modified Exact test, BH-corrected *P* = 1.3 × 10^-8^), demonstrating that genetic disruption of a wide variety of biological processes leads to severe proliferation phenotypes in hNSCs.

To gain insight into the biology associated with changes in hNSC proliferation we utilized a public database of manually curated and peer-reviewed pathways to test the enrichment of gene proliferation phenotypes within signaling pathways (table S5) (*42*). We found that gene disruption phenotypes are significantly enriched for the FGF signaling pathway (hypergeometric test, BH-corrected *P* = 2.8 × 10^-2^) (Fig. 2D; table S5) consistent with its role in anterior forebrain cortical patterning (*44, 45*). Gene disruption phenotypes are also enriched for the WNT signaling pathway (hypergeometric test, BH-corrected *P* = 2.1 × 10^-2^) (table S5), which contributes to cortical patterning via a signaling center in the cortical hem (*46*). In addition, we observe enrichment for ROBO signaling (hypergeometric test, BH-corrected *P* = 3.1 × 10^-7^) (table S5) supporting evidence from mammalian genetic models that this pathway alters hNSC self-renewal (*47*). During human corticogenesis, neural progenitors are influenced by signaling molecules released from nearby patterning centers and these morphogenetic gradients result in the specification of neuronal cell types and cortical areal identity (*46*). Genetic disruption impacting these signaling pathways may alter neurogenesis and possibly result in changes to the specification of cortical size and areal identities (*45, 46*).

To explore genetic disruptions altering enhancer function, we used PCA to isolate the magnitude and temporal effects of disruption for all conserved enhancer regions included in our screen (Fig. 3A; table S6). We combined this analysis with genome-wide predictions of transcription factor binding sites (TFBS) to identify binding sites enriched in proliferation-altering enhancer disruptions. Most conserved regions included in our screen (90.7% of total conserved regions) include a predicted TFBS, with a total of 152,110 motifs predicted across all targeted regions (*49*). To obtain an initial view of the effect of genetic disruption on predicted TFBS motifs, we individually interrogated 8 conserved regions targeted in our screen, including a subset exhibiting proliferation phenotypes. We performed high-throughput amplicon sequencing on the targeted conserved regions after genetic disruption in order to determine the molecular effects on enhancer TFBS motif content (fig. S9-10; table S7). In most cases (87.5% of replicated sgRNAs) we observed genetic variation at the sgRNA-Cas9 target site. The proportion of alleles modified following disruption ranged from 33-96% (table S7) and deletions were the most common type of genetic variation observed (average deletion size of 6 to 7 bp). The 8 individually targeted conserved regions overlap 50 predicted TFBS motifs, and 41 motifs were likely disrupted due to proximity (within 10 bp) to the predicted sgRNA-Cas9 cleavage site. One proliferating-decreasing disruption destroyed a putative TBX2/TBX20 TFBS motif that includes human-specific substitutions within *HACNS96* (Fig. 3A,C; fig. S9). TBX2 mediates regulation of FGF signaling during anterior neural cell specification (*50*). Transgenic assays demonstrate that *HACNS96*, which is located within the intron of the transcription factor *NPAS3*, acts as a transcriptional enhancer during vertebrate neurodevelopment (*41*). *NPAS3* is expressed in the developing brain and genetic disruption of *NPAS3* results in a proliferation-decreasing phenotype (Fig. 2A), suggesting that disruption of the TBX2/TBX20 motif within *HACNS96* may lead to the proliferation-decreasing phenotype we observed by altering *NPAS3* regulation.

**Fig. 3.**
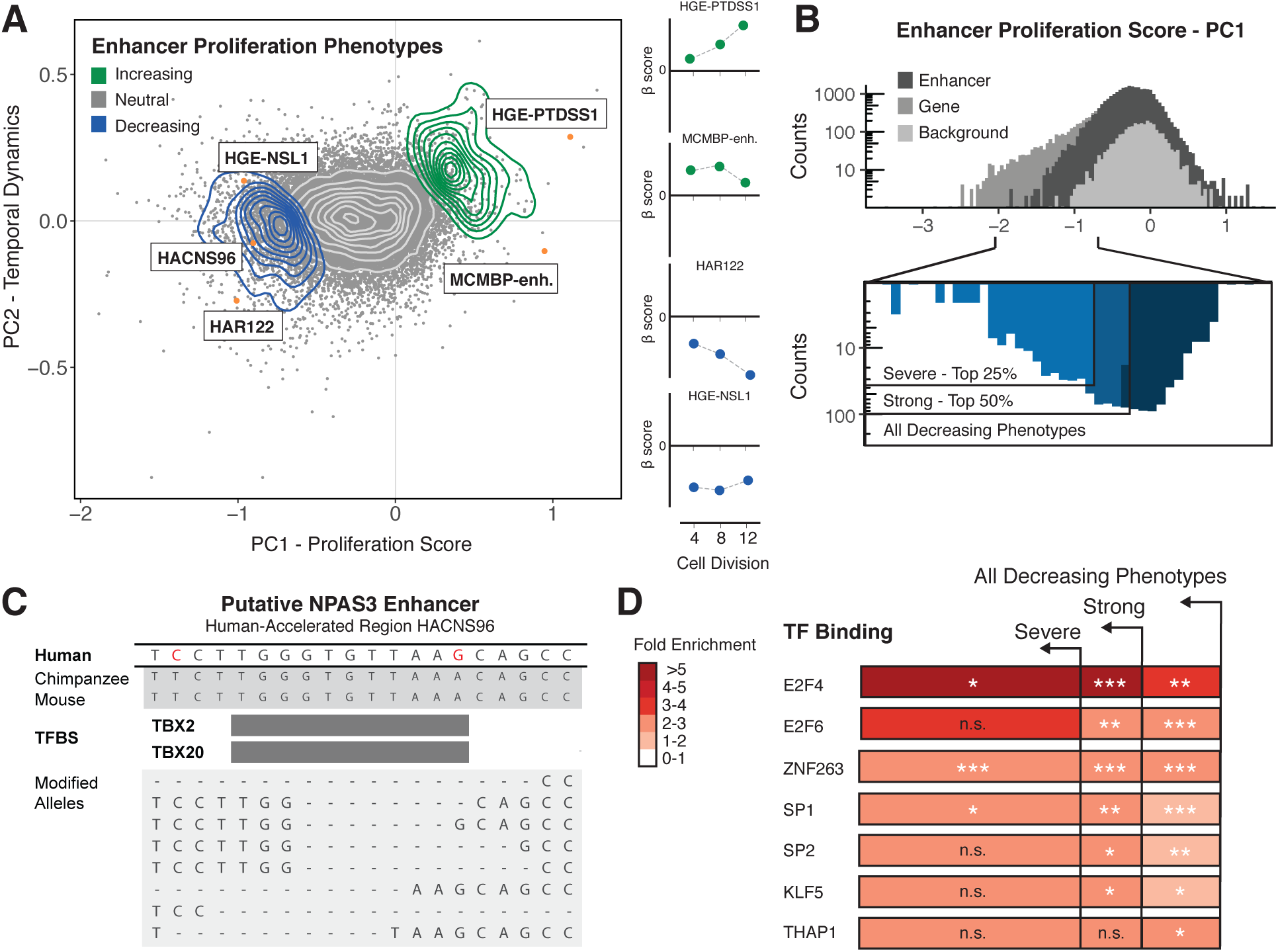
Characterization of enhancer proliferation phenotypes. (**A**) (*Left*) Principal component analysis (PCA) of all gene disruptions with a 2-D density overlay illustrating the distribution of proliferation-decreasing (*blue*), neutral (*grey*), and proliferation-increasing (*green*) phenotypes. Individual conserved region disruptions (*orange*) with biological roles in the maintenance of neural stem cells or human neurodevelopment (*6, 34, 41, 48*). (*Right)* Temporal dynamics captured by principal component analysis shown at 4, 8, and 12 cell divisions. (**B**) (*Top*) Histogram of proliferation scores obtained from PCA for enhancer disruptions (*black*) and genomic background controls (*grey*). (*Bottom*) Partitioning of all enhancer disruptions that decrease proliferation into severe (top 25%) and strong (top 50%) categories. **(C)** (*Top*) Genomic alignments for human, chimpanzee, and mouse for a conserved region within a HAR enhancer (*HACNS96*). Human-specific substitutions (*red*) indicate genetic changes occurring within the conserved regions targeted for genetic disruption in the screen. (*Bottom*) Examples of deletion alleles introduced by Cas9 in this locus, determined by long read amplicon resequencing and alignment to the reference genome (GRCh37/hg19). (**D**) Predicted transcription factor binding sites were obtained from the JASPAR 2018 TFBS prediction database (*49*) significantly enriched in proliferation-altering enhancer disruptions within the partitions shown in **(B)**. n.s. = not significant, * = hypergeometric BH-corrected p-value < 0.05; ** = hypergeometric BH-corrected p-value < 0.005; *** = hypergeometric BH-corrected p-value < 0.0005.

To identify predicted TFBS motifs that are overrepresented in proliferation-altering enhancer disruptions, we partitioned the proliferation-decreasing phenotypes within enhancers into severe, strong, and all decreasing categories (Fig. 3B). Conserved regions exhibiting severe proliferation-decreasing phenotypes are enriched in E2F4 and E2F6 binding motifs (hypergeometric test, BH-corrected *P* = 2.6 × 10^-2^ and *P* = 5.9 × 10^-4^, respectively) (Fig. 3D), consistent with the role of E2F factors in the transcriptional control of cell cycle dynamics and cell specification (*51*). We also observed enrichment of ZNF263, SP1, and SP2 (hypergeometric test, BH-corrected *P* = 6.8 × 10^-12^, *P* = 6.3 × 10^-4^, *P* = 4.7 × 10^-3^, respectively) indicating that binding of broadly expressed general transcription factors is important in facilitating normal enhancer function. We did not observe enrichment for transcription factor binding sites within proliferation-increasing disruptions, potentially due to the smaller number of conserved regions involved in this cellular phenotype.

To describe the proliferation-altering phenotypes at the level of whole enhancers, we summarized the number of conserved region disruptions impacting proliferation within each targeted enhancer (table S9). While many enhancers include only one disrupted site that results in a proliferation phenotype (66.9% of proliferation-altering enhancers), we also identified many enhancers that included multiple conserved regions impacting proliferation (Fig. 4A). On average, 15% of conserved regions within proliferation-altering enhancers were associated with a proliferation phenotype (Fig. 4B). The cumulative burden of proliferation-altering disruptions within whole enhancers scales linearly with the number of targeted sites (Fig. 4C), supporting that the total proliferation phenotype burden is proportional to the number of conserved regions disrupted within each enhancer.

**Fig. 4.**
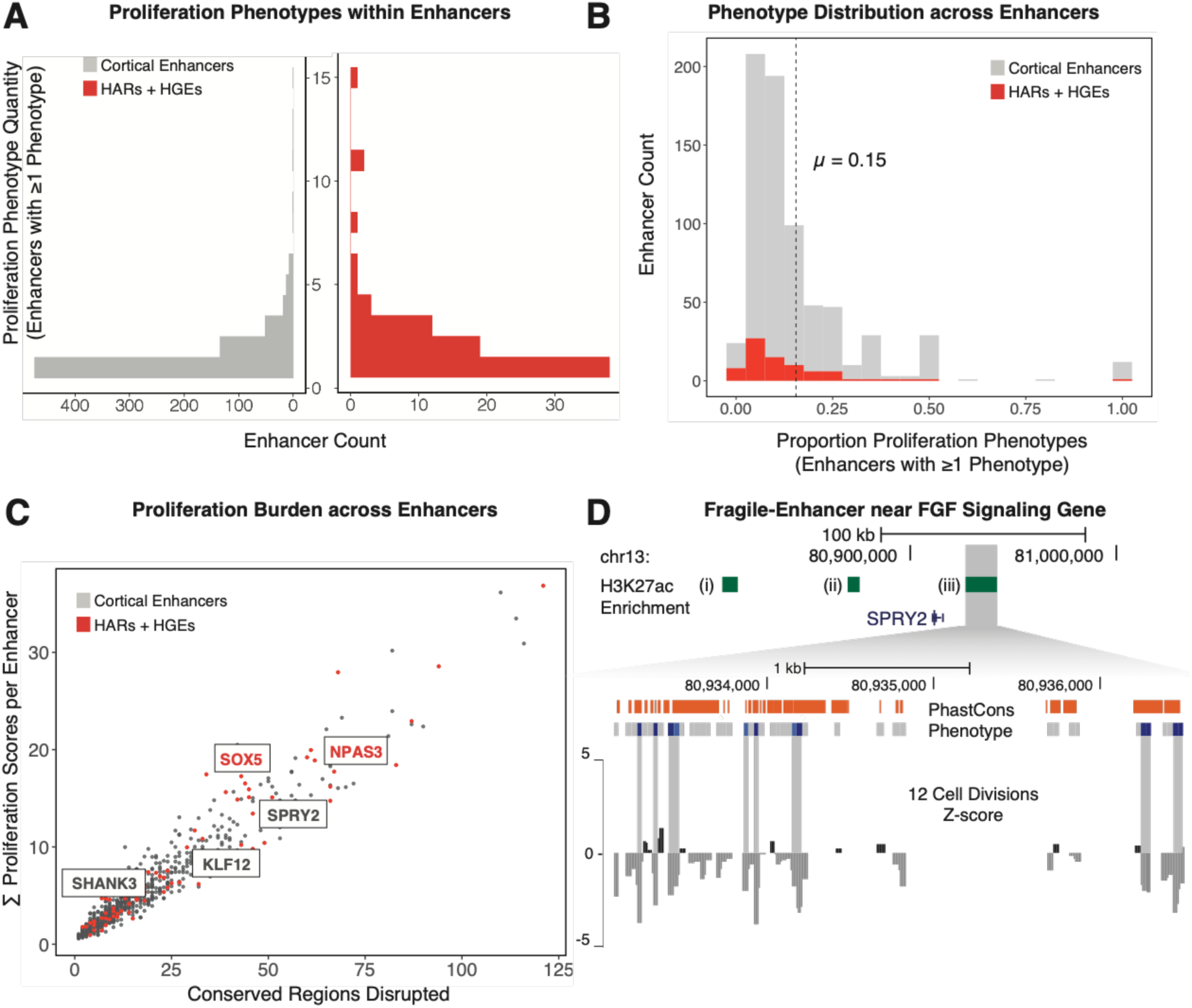
Frequency of proliferation-altering disruptions in enhancers, HARs and HGEs. (**A**) The number of disrupted conserved regions leading to a proliferation phenotype within each enhancer (left, shown in grey) or in each HAR or HGE (right, shown in red). (**B**) The proportion of proliferation-altering disruptions compared to all targeted sites in all enhancers (*grey*) and HARs or HGEs (*red*). The dashed line indicates the mean density of proliferation phenotypes within all enhancers in the screen. (**C**) The total number of conserved regions disrupted within each enhancer (*x-axis*) compared to the cumulative proliferation phenotype burden within each enhancer (*y-axis*). Fragile enhancers identified by permutation analysis (see panel **D** and main text for details) are labeled based on the most proximal gene. HARs and HGEs are labeled in red. (**D**) A fragile enhancer near *SPRY2*, a regulator of the FGF signaling pathway. (*Top)* Genomic coordinates indicate position on the reference genome (GRCh37/hg19). Three enhancers (i, ii, iii), shown by the extent of H3K27ac marking (*green bars*) in hNSCs, were disrupted in this locus. Enhancer (iii) includes a significant excess of proliferation-altering enhancer disruptions. (*Bottom)* Mammalian PhastCons elements (*orange bars*) and corresponding conserved regions that have either neutral effects (*grey bars*) or proliferation-decreasing phenotypes (*blue bars*) when disrupted in the screen are shown. The signal track at the bottom of the figure displays normalized beta scores (in grey) at 12 cell divisions averaged across a sliding window.

We then identified regulatory elements that we termed “fragile enhancers”, which are a subset of enhancers found by permutation analysis to contain a significant excess of conserved regions yielding a proliferation phenotype (supplementary text). These fragile enhancers are exceptionally sensitive to genetic disruption, and variation within them may have substantial effects on human cortical development (table S9). One fragile enhancer (permutation test, BH-corrected *P* = 7.4 × 10^-3^) is proximal to *SPRY2* (Fig. 4D), a known regulator of FGF signaling (*52*), illustrating the potential vulnerability of hNSCs to variation influencing this developmental signaling pathway. We identified two fragile enhancers that include the human-accelerated regions *HACNS610* (permutation test, BH-corrected *P* = 4.4 × 10^-3^) and *HAR122* (permutation test, BH-corrected *P* = 4.4 × 10^-3^) located within introns of *SOX5* and *NPAS3,* respectively. The identification of fragile enhancers containing HARs suggests candidates for human-specific regulatory activity at important positions within regulatory networks impacting hNSC self-renewal.

To link enhancers exhibiting proliferation phenotypes with their putative target genes, we used a high-resolution map of long-range chromatin contacts ascertained from human neural precursor cells (table S10; supplementary text) (*53*). Chromatin interaction maps (Fig. 5A) identified 180 enhancer-gene pairs (table S10) between enhancers and genes that exhibit proliferation phenotypes in hNSCs. These maps capture a diverse range of regulatory interactions, including interactions between a single enhancer and a single gene target; interactions between multiple enhancers converging on a single gene target; and a single enhancer that interacts with multiple target genes. Likely because of this diversity, we did not observe a clear correlation (Pearson correlation = 0.01) between the effect of enhancer and target gene disruptions. However, individual enhancer-gene interactions provided insight into the relative effects of enhancer versus target gene disruption on proliferation.

**Figure 5.**
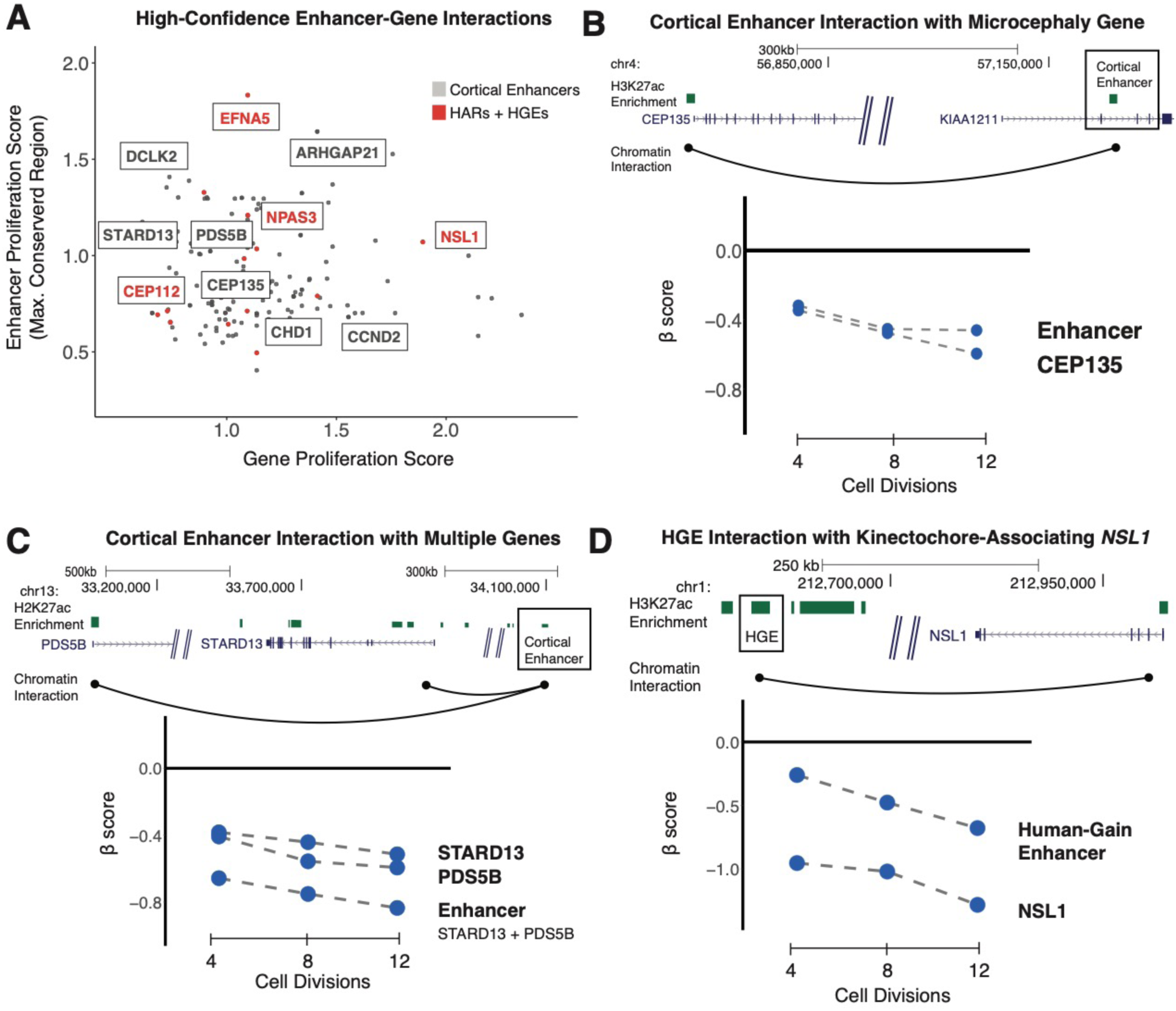
Identifying target genes of enhancers contributing to hNSC proliferation using chromatin contact maps. (**A**) Chromatin contact maps within human neural precursor cells (*53*) were used to identify interactions between enhancers and genes that, when disrupted, result in a proliferation phenotype. For each enhancer-gene pair, the absolute value of the gene proliferation score (*x-axis*) and the absolute value of the strongest proliferation score within the enhancer (*y-axis*) is shown. Highlighted genes have known roles in neural stem cell biology or are associated with risk for developmental disorders (*31, 48, 55*). Interactions between HARs or HGEs and target genes are labeled in red. (**B**) An example of an enhancer (boxed) regulating a single target gene associated with microcephaly (*31*). (*Top*) The genomic coordinates (in GRCh37/hg19) of the enhancer are shown relative to the two detected target genes. Chromatin contacts are shown as black arcs. (*Bottom*) The effects of enhancer and target gene disruptions on hNSC proliferation at 4, 8 and 12 cell divisions. (**C**) An example of an enhancer (boxed) regulating multiple target genes with roles in hNSC proliferation. The figure is labeled as in **B.** (**D**) An example of a human-gain enhancer (HGE, boxed) interacting with the nearby gene *NSL1*. The figure is labeled as in **B.**

The microcephaly-associated gene *CEP135* (*31*) interacts with a single enhancer active during human corticogenesis (Fig. 5B). Disruption of the enhancer and of *CEP135* result in comparable perturbation of hNSC proliferation, suggesting that cortical phenotypes may arise through changes in the regulation of neurodevelopmental risk genes. We also identified a human cortical enhancer (Fig. 5C) interacting with two target genes with known roles in cell proliferation, *STARD13* and *PDS5B* (*54, 55*). Genetic disruption of the enhancer results in a stronger effect on proliferation than disruption of either target gene (Fig. 5C, bottom), suggesting that genetic variation within a single enhancer can lead to severe biological effects through dysregulation of multiple genes.

We also utilized this chromatin interaction map to identify gene targets for 9 HARs and 8 HGEs that contribute to hNSC proliferation. One example, shown in Figure 5C, is an HGE that targets *NSL1*, which functions in neural progenitor cell division and has been associated with cognitive phenotypes in humans (*48, 56*). Disruption of *NSL1* negatively affects hNSC proliferation (Fig. 5D, bottom). Disruption of the interacting HGE also alters proliferation, resulting in a weaker, but significant biological effect compared to the gene disruption. These findings provide the basis for determining whether gains in activity in HARs and HGEs alter expression of specific gene targets during neurogenesis.

Our results revealed genes and enhancers that regulate hNSC proliferation and that, when disrupted, potentially alter cortical development. Genes associated with developmental disorders, including autism spectrum disorder, disproportionately impacted proliferation, suggesting one mechanism contributing to the etiology of those phenotypes. Disruptions in many enhancers active during human corticogenesis also led to alterations in hNSC proliferation. These enhancers include HARs and HGEs, providing clear evidence that regulatory elements linked to human brain evolution play an important functional role in neurogenesis. TFBS motifs enriched in proliferation-altering regions highlight the importance of specific transcription factors, and individual binding sites, in transcriptional regulation within the neural stem cell niche. Integrating enhancer and gene proliferation phenotypes with enhancer-gene interactions revealed the diverse ways in which the biological effects of enhancer disruptions manifest via their target genes. Collectively, our findings provide a basis for interpreting genetic variation within regulatory elements and identifying enhancers with central roles in human cortical development, neurodevelopmental disorders, and human cortical evolution.

## Acknowledgements

We thank R. Muhle for comments on the manuscript; S. Mane, K. Bilguvar, and C. Castaldi at the Yale Center for Genome Analysis for sequencing data; B.Fontes and R. Airo at Yale Environmental Health and Safety for supporting the lentiviral work; and the members of the PsychENCODE Consortium for providing chromosome conformation capture data to the research community

## Author Contributions

E.G., D.E., J.C., and J.P.N. conceived and designed the study. E.G. designed and implemented the pipeline for sgRNA selection within conserved regions. J.G. advised the construction of the sgRNA-Cas9-GFP plasmid libraries. E.G. generated the sgRNA-Cas9-GFP plasmid libraries for proliferation screening experiments. E.G. generated the lentiviral libraries and conducted the human neural stem cell transduction experiments. E.G. designed and implemented the proliferation phenotype analysis pipeline. E.G. performed TF binding site and permutation-based enhancer proliferation phenotype enrichment analyses. S.U. provided statistical advice for enrichment analyses. E.G. generated constructs for amplicon resequencing experiments. E.G. and J.P.N. wrote the paper with input from all authors.

## Data and Materials Availability

Sequencing data for RNA-seq and H3K27ac enrichment collected from H9-dervived human neural stem cells are available through the Gene Expression Omnibus under accession number GSE57369. Chromosome conformation capture data for human neural progenitor cells are available through the PsychENCODE Knowledge Portal hosted under the Synapse ID syn12984505. Sequencing data for massively parallel sgRNA-Cas9 disruption and individual replicate disruption are available through the Gene Expression Omnibus under accession number GSE138823. All code used in the analysis is available on GitHub (https://github.com/NoonanLab/Geller_et_al_Enhancer_Screen).

## Materials and Methods

### Cell Lines and Vectors

Materials were obtained from the following sources: H9-derived human neural stem cells from Life Technologies (N7800-1000), HEK293FT cells from Invitrogen, LentiCRISPRv2GFP from Addgene (Plasmid #82416, provided by David Feldser), pCMV-VSV-G (Addgene, Plasmid #8454), and pCMV-dR8.2 dvpr (Addgene, Plasmid #8455). The identity of H9-derived human neural stem cells was confirmed by analysis of hNSC markers via RNA-sequencing and antibody staining for multipotency markers (SOX2, NES). The manufacturer provides protocols for maintaining hNSC and differentiating hNSC into neurons (MAP2, Dcx), oligodendrocytes (GalC, A2B5), or astrocytes (CD44, GFAP) (*57*).

### Human Neural Stem Cell Culture

H9-derived human neural stem cells were cultured in Knockout DMEM/F-12 (Life Technologies, N7800-100) supplemented with EGF anwd FGF (ConnStem), GlutaMax-I and StemPro neural supplement (Life Technologies) as recommended by the manufacturer. In addition, cells were cultured on Matrigel–coated flasks seeded at ∼50,000 cells per square centimeter. The doubling time of H9-derived human neural stem cells is approximately 48 hours (*57*).

### K-Means Clustering for Identification of Cortical-Enriched Enhancers

H3K27ac data from human cortex was compared with publicly available H3K27ac datasets as follows: A single composite multi-sample enhancer annotation from developing cortex, limb, embryonic stem cells, and select adult tissues profiled by the Roadmap Epigenomics Project (*58*) was generated by merging replicate peaks across all samples using a 1bp overlap. The level of H3K27ac signal in each region for each sample was quantified by averaging read counts per kilobase per million mapped reads (RPKM) in each region from each replicate. Each region was represented by a vector of a length equal to the total number of tissues considered, with each point representing the RPKM value of marking in that region for a single tissue. Each vector was normalized by subtracting the mean of all tissue quantifications from each individual tissue quantification, divided by the standard deviation of values for that vector. The matrix of these normalized tissue quantification values was then subjected to k-means clustering using R to identify sets of sites exhibiting the strongest marking in each tissue compared to all other samples in the comparison. We used GREAT version 3.0.0 (http://great.stanford.edu/) to identify biological functions and processes showing significant enrichment for each set of enhancers (*59*).

### sgRNA Library Design: Defining Genomic Background Controls

A set of background controls were defined by initially shuffling the location of randomly selected subset of targeted enhancers (n=500). Next, the PhastCons elements underlying the original enhancers were shuffled within the shuffled enhancers; these pseudo-PhastCons elements residing in shuffled enhancers were termed ‘genomic background controls.’ Individual sgRNAs targeting the genomic background controls were scored and filtered in the same manner as the enhancers described above. In addition, these regions were filtered for possible regulatory function based on evidence of epigenetic activity across stem cell and brain tissue-types. Specifically, DNase-hypersensitive sites (DHSs) identified in H9-derived neural progenitor cells were extended by 1,000 bp (*60*); genomic background control regions overlapping the extended DHSs by 1 bp were excluded from downstream analyses. In addition, a compendium of epigenetic atlases (Epilogos, https://epilogos.altius.org) was utilized to filter remaining genomic background controls for evidence of gene regulatory function based on chromatin state across a variety of human tissue-and cell-types (*61*). Chromatin states across ‘All 127 Roadmap Epigenomes’, ‘Brain’, and ‘ESC derived’ were used to filter genomic background controls based on cumulative evidence across the shuffled enhancers. The following criteria was applied to filter shuffled enhancers: evidence across all chromatin states (not including ‘Quiescent’ states) was set to ‘All 127 Roadmap Epigenomes’ < 5.0, ‘Brain’ < 0.5, and ‘ESC derived’ < 0.5. Genomic background regions within shuffled enhancers passing the filtering criteria for DHSs and chromatin state (described above) were included in downstream analyses.

### sgRNA Library Design: Defining Proliferation-Decreasing Genes

Proliferation-decreasing controls were identified from Wang, et al. (2015) as sgRNAs exhibiting essential gene scores across a panel of cancer cell lines including KBM7, K562, Jiyoye, and Raji (*14*). Genes identified as proliferation-decreasing control targets exhibit a CRISPR-score (‘CS’) < −2.0 and ‘adjusted p-value’ < 0.05 across all 4 cell lines performed by Wang et al. (2015). Individual sgRNAs (up to 10 sgRNAs per gene) targeting genes that meet these criteria were used as a control for proliferation-decreasing genes in this study.

### sgRNA Library Design: Scoring and Filtering sgRNAs

For enhancer regions, sgRNAs were designed and scored across PhastCons elements (Placental Mammal Conserved Elements) (*24*). For protein-coding regions, sgRNA designs were included from Wang et al. (2015) (*62*). For enhancer, gene, and background control (see below) sgRNAs, the scoring metric was incorporated from Gilbert et al. (2014) (*63*). Bowtie version 1.1.2 was used to score mismatches across the genome (version GRCh37/hg19) (*64*); a score of e29m1 was used as a cutoff for potential sgRNAs. The scoring scheme is summarized as follows: the sgRNA sequence is extended to a 23-mer including the PAM motif NGG. Genome-wide mapping with Bowtie was used to score each sgRNA based on the matrix [9,9,9,9,9,9,9,9,19,19,19,19,19,28,28,28,28,28,28,28,0,19,40] where the PAM motif is represented at the end of the matrix. Then, using the filtering criteria -e29m1 allows sgRNAs with mismatches unlikely to result in cleavage, then excludes sgRNAs if more than 1 genome mapping event is reported. All sgRNAs were then filtered based on GC-content 20-80% and excluding poly-T sequences greater than 4 in length.

### sgRNA Library Design: Defining Non-Targeting Control sgRNAs

Non-targeting controls (n=500) were generated from random sgRNA sequences that were processed by the same scoring procedure as enhancer regions described above, including filtering based on GC-content 20-80% and poly-T sequences greater than 4 in length. As with enhancer-targeting sgRNAs, Bowtie was used to filter sgRNAs based on the scoring matrix [10,10,10,10,10,10,10,10,10,10,10,10,10,10,10,10,10,10,10,10,0,19,40] where the PAM motif is represented at the end of the matrix. The following mapping criteria were used: -e39m1, --max, and –un; sgRNAs that yield no mapping with up to 3 mismatches permitted across the reference genome (GRCh37/hg19) were reported as unmapped; this subset of sgRNAs was included as non-targeting controls.

### sgRNA Library Design: Selecting sgRNAs Targeting Enhancers and Genomic Background Controls

To select sgRNAs, an R script processed filtered sgRNAs with the following procedure: each targeted PhastCons element was extended by 15 bp and the extended PhastCons element was partitioned into 30bp windows. Then, sgRNAs were randomly drawn with uniform probability from the filtered sgRNA set for each window such that up to 2 unique sgRNAs were selected per window, and at least 2 sgRNAs were selected per PhastCons element. Gencode (v19) (*65*) was utilized to exclude conserved regions overlapping gene promoters (+/-1 kb from TSS) and exons for all coding transcripts with evidence of level 1 or 2 support (validated or manual annotation).

### sgRNA Library Design: Selecting sgRNAs Targeting Protein-Coding Regions

For protein-coding regions, genes were selected for targeting based on expression levels measured by RNA-sequencing in two biological replicates of H9-derived human neural stem cells (FPKM > 1 across two replicates) (*18*); this yielded 10,674 expressed genes. All protein-coding sgRNAs from Wang et al. (2015) (*62*) were processed through the scoring and filtering procedure described above. Next, two filtered protein-coding sgRNAs were randomly selected for each gene.

### sgRNA Library Design: Specificity Controls

To assess the proportion of on-target activity resulting from sgRNAs with mismatches in the targeting sequence, we generated a set of specificity controls. Gene targeting sgRNAs for *CCND2 SOX2*, and *SRSF1* were included as a reference for on-target activity. The 20 nt PAM-adjacent targeting sequence was divided into 3 regions based on the tolerance of Cas9 to mismatches (Region 1 is the PAM-adjacent region (1-7 nt), Region 2 (8-12 nt), and Region 3 (13-20 nt) is distal to the PAM-adjacent region) (*63*). To determine the sensitivity of on-target activity to mismatches (MM), single nucleotide mismatches were introduced into the target sequence within each region or spanning multiple regions and the number of mismatches ranged from 1 MM to 4MM.

### sgRNA Library Design: Assigning Sub-Libraries for Genetic Screening

All sgRNAs were divided across 8 sub-libraries (‘subpools’) for large-scale transduction into hNSCs (6 enhancer-targeting subpools and 2 gene-targeting subpools). For each enhancer targeted by the screen, the enhancer was randomly assigned to one of the six subpools and all sgRNAs targeting that enhancer were assigned to the same subpool. For each gene targeted by the screen, the gene was assigned to one of the two subpools and all sgRNAs targeting that gene were assigned to the same subpool. In addition, each subpool included identical sets of non-targeting control sgRNAs (described above) and proliferation-decreasing sgRNAs identified in previous sgRNA-Cas9 genetic screens (*62*).

### Lentiviral sgRNA Plasmid Library Construction

Oligonucleotides were synthesized on a CustomArray 90K array (CustomArray, Inc). The first round of PCR amplified sub-library specific sgRNA sequences (S01-S08). The second round of PCR introduced overhangs compatible for Gibson assembly (New England Biolabs) into LentiCRISPRv2GFP linearized with BsmBI. PCR reactions were monitored using SYBR green to ensure each reaction was terminated in the linear amplification phase. Gibson Assembly reaction products were purified and transformed into E. Coli DH10B (Life Technologies). To preserve the diversity of the library, at least 500-fold coverage in library representation was recovered in each transformation, and each transformed library was grown in liquid culture until OD 0.8-1.0 (∼8 hours).

PCR primers for sub-library amplification:

S01

F-ACCCAAAGAACTCGATTCCT

R-ATGGAGGTCCTTTTGTTCCT

S02

F-AGCGTCGAATGAATGCATAC

R-AACTTCAGGGCTGTGTCTAA

S03

F-AGACCAGGATGGCTGATAAG

R-GTTTCGTGCCCACATATACC

S04

F-AATCCTTGCGTCAATGGTTC

R-GGGTTCTCGGATTTTACACG

S05

F-TGTCGTGCCTCTTTATCTGT

R-GCTTCGGTGTATCGGAAATG

S06

F-TTCCGTTTATGCTTTCCAGC

R-TCCTTGGAGTTTAGAGCGAG

S07

F-TGCAAGTGTACAAATCCAGC

R-GAACGGTGATCCCTTTCCTA

S08

F-TTATAATCATCCTCCCCGGC

R-CCAAATAGGATGTGTGCTCG

PCR primers for Gibson assembly homology arms:

F-GGCTTTATATATCTTGTGGAAAGGACGAAACACCG

R-ACTTTTTCAAGTTGATAACGGACTAGCCTTATTTTAACTTGCTATTTCTAGCTCTA AAAC

Individual sgRNA representation within each plasmid library (S01-S08) was determined by high-throughput 2×100 bp sequencing on the HiSeq4000 instrument (Illumina) (Fig. S2).

### Lentivirus Production

Lentiviral work was performed using BSL-2 Plus safety procedures, including production of lentiviral sgRNA-Cas9 libraries, culturing of transduced cells, and extraction of genomic DNA. Lentivirus was produced by co-transfecting the sgRNA-Cas9-GFP library vector with pCMV-VSV-G and pCMV-dR8.2 dpvr packaging plasmids into HEK293FT cells using Extreme Gene 9 transfection reagent (Millipore-Sigma) in serum-free media supplemented with GlutaMax-I (ThermoFisher) and 25uM chloroquine (Millipore-Sigma). After 8 hours, media was replaced with high bovine serum albumin (Millipore-Sigma) (1.1g/100mL) in GlutaMax-supplemented OptiMem (ThermoFisher) with 10uM sodium butyrate (Millipore-Sigma). The virus-containing supernatant was collected 48 hours after replacement. Viral supernatant was passed through a 0.45uM low-binding filter and immediately concentrated using Amicon Ultra-15 100kD filters. Concentrated virus was aliquoted, flash-frozen over dry ice and stored at −80C.

### Large-Scale Human Neural Stem Cell Transduction

Target cells in 25 cm tissue culture flasks at 250,000 cells per sq cm were transduced in low volume media containing 8ug/mL polybrene (Millipore-Sigma); 24 hours after infection virus was removed and cells were passaged to a density of 50,000 cells per sq cm. To establish lentiviral titer, serial dilutions of concentrated virus were added to 25 cm tissue culture flasks and incubated for 24 hours. Cells were then passaged to a density of 50,000 cells per sq cm and infection rate was determined 48 hours later using GFP expression measured by flow cytometry (BD Accuri C6). For high-throughput screening, each sub-library was transduced by plating 50 million cells across eight 25 cm flasks and adding the appropriate volume of lentivirus to each flask. Initial multiplicity-of-infection was ∼0.3-0.4 to achieve >1000-fold library coverage and infection was monitored after 24 hours by flow cytometry for GFP expression over the course of the experiment (at 4, 8, and 12 cell divisions). Cells were harvested at each passage and stored as a cell pellet at −20C for genomic DNA extraction.

### sgRNA Library Readout by High-Throughput Sequencing

Each sgRNA subpool library readout was performed using two steps of PCR as described in Chen et al. (2015) (*64*). Second round PCR products were purified using column-based cleanup (New England Biolabs). Second round PCR products containing Illumina adapters at each timepoint belonging to a single subpool (e.g., S01) and biological replicate were combined and submitted for sequencing on the same channel(s) of a single sequencing run. Diluted libraries were spiked in with whole-exome libraries and sequenced using 2×100 bp reads on the HiSeq 4000 system (Illumina).

### Mutation Spectrum of Individual Conserved Region sgRNAs

Individual sgRNA-Cas9-GFP plasmids were cloned, propagated in Stable Competent E. Coli. strains (NEB), and isolated using Endo-Free Maxi Prep Isolation Kits (Thermo-Fisher). Transient transfection of 4 million hNSC per construct was achieved using the Mouse Neural Stem Cell Nucleofection Kit (Amaxa). At 96 hours post-transfection, GFP-positive cells were separated on an S3e Cell Sorter (BioRad) followed by DNA extraction. Amplicons from individual sgRNA-mediated were analyzed by high-throughput sequencing followed by insertion, deletion, and substitution analysis using CRISPResso2 (Fig. S9-S10) (*66*).

### Identification of Proliferation Phenotypes

To quantify the biological effects of disruption on cell proliferation, we utilized a model-based analysis of genome-wide CRISPR-Cas9 knockout (MAGeCK) which models read counts using the negative binomial distribution (*67*). First, each sgRNA is assigned to a target representing either a gene or conserved region. Each target can contain one or more sgRNAs that can be jointly modeled in the MAGeCK analytical pipeline. Sequencing reads were initially filtered using CutAdapt version 1.16 and options (-j 20, -l 20, -g GACGAAACACCG, --trimmed only). Trimmed reads were used as input into MAGeCK version 0.5.8 which was performed for replicates individually and jointly. The following options were used: --norm-method control, --control-sgrna NTC_sgRNA_ID.txt. MAGeCK analysis was conducted for each sub-library independently, and the same panel of non-targeting controls was included in each sub-library and used to normalize read counts. The results of each sub-library were combined using custom R scripts. For the replication plots (Fig. 1B-C), MAGeCK was performed independently for each replicate. For all subsequent analysis and values reported in tables, the results are from jointly modeling the biological effects across two replicates. MAGeCK provided an estimate of the biological effect following genetic disruption on cellular proliferation termed the β score. For each conserved region, the β score is associated with a permutation-based p-value determined by permuting sgRNAs targeting each conserved region and evaluating the probability of observing the biological effect within the set of genomic background control sgRNAs.

Cell proliferation phenotypes were identified using linear-discriminant analysis (LDA) on a training set then applying the learned classifier to the full dataset (*68*). Similar approaches have been used to characterize phenotypes following high-throughput editing experiments (*26*). LDA produces a classification that maximizes the separability of the input groups. The training set was defined as follows: the ‘decreasing’ population includes 500 genes decreasing proliferation across a panel of cancer cell lines (Wang, et al. 2015) (*62*), the ‘neutral’ population is all regions within the genomic background, and the ‘increasing’ population is comprised of the top 1% of proliferation increasing regions identified at each time point (4, 8, and 12 cell divisions) (Fig. S7). All sgRNAs for the training set were included in the hNSC proliferation screening experiments. LDA was performed jointly across timepoints, and each disrupted region (gene and conserved regions within enhancers), was classified based on a composite of the regression-based effect size and p-value.

A composite score was then obtained by multiplying the MAGeCK β score by the permutation-based p-value for phenotype classification using LDA. The resulting proliferation phenotype classifications (‘decreasing’, ‘neutral’, or ‘increasing’) were filtered for reproducible effect sign across biological replicates and consistent phenotype classification across 4, 8, and 12 cell divisions (e.g. proliferation-decreasing at 4, 8 and 12 cell divisions).

MAGeCK β scores and proliferation phenotype classifications at 4, 8, and 12 cell divisions are available for visualization in the UCSC Genome Browser (GRCh37/hg19): http://noonan.ycga.yale.edu/noonan_public/Geller_Enhancer_Screen/hub.txt

### Principal Component Analysis and Pearson Correlation Analysis

To extract latent factors and perform correlation analyses, we constructed a single composite annotation of all β scores for each ‘subpool’ across multiple timepoints. These Beta scores were assembled into a single data matrix using custom R scripts. Each row in this matrix represented a single conserved region or gene disruption, and each column represented the Beta score at 4, 8, or 12 cell divisions. Principal component analysis was performed on this matrix using R (Fig. 2A; Fig. 3A). Pearson correlation analysis was also carried out using R (Fig. S6).

### Transcription Factor Binding Site Enrichment Analysis

We collected 572 transcription factor binding site (TFBS) predictions from the JASPAR 2018 database (*49*) that overlap with at least one conserved region included in this study. To identify TFBS that are enriched within conserved regions that have biological effects on proliferation, we conducted hypergeometric tests using custom R scripts. The hypergeometric test was conducted for each TFBS independently. Each test was constructed to compare the abundance of TFBS in the category of interest (‘severe’, ‘strong’, ‘all’, or ‘positive’) and compared to ‘neutral’ phenotype conserved regions. The hypergeometric p-value for assessing the enrichment of each TFBS for all sites were then corrected for multiple-testing using the Benjamini-Hochberg procedure (*69*).

### Enhancer Proliferation Phenotype Permutation Analysis

We used permutation analysis to identify enhancers containing a significant excess of proliferation phenotypes. The procedure was implemented using custom R scripts. For each permutation, proliferation phenotypes were randomly shuffled across all conserved regions. We performed 100,000 permutations. The significance of proliferation phenotypes within an enhancer was assessed based on the fraction of permutations where the number of proliferation phenotypes was greater than or equal to the number of proliferation phenotypes observed within the enhancer. The resulting permutation-based p-values were then corrected for multiple-testing using the Benjamini-Hochberg procedure (*69*).

### Identification of Enhancer-Gene Interactions

Hi-C data from human neural precursor cells were generated by the PsychEncode Consortium (*70*). High-confidence loop calls and 50-kb topologically associating domains (TADs) were made available by the authors through the Synapse repository. The Juicer tool suite was utilized to identify contact domains using the default settings and the ‘arrowhead’ algorithm (*71*). Custom R scripts implemented the following procedure to generate enhancer-gene interactions. Enhancers were defined as regions containing at least one proliferation phenotype. Gencode (v19) (*65*) was utilized to define gene regions including the promoter (+10 kb), transcription start site, and gene body. Genes harboring a proliferation phenotype were used to identify enhancer-gene interactions. For loop calls, anchor points were used to identify enhancer-gene interactions. For contact domains, enhancers were associated with each gene harboring a phenotype within the contact domain. High-confidence interactions were reported as enhancer-gene interactions derived from loop calls and contact domains. In addition, topologically associated domains (TADs) were used to identify enhancer-gene interactions. For TADs, enhancers with at least one phenotype within the TAD were associated with all genes harboring proliferation phenotypes within the TAD.

**Table S1.**

Sample statistics for each sub-library and transduction replicate in the study.

**Table S2.**

Results from proliferation analysis following gene disruption.

**Table S3.**

Results from proliferation analysis following gene disruption for developmental disorder risk loci.

**Table S4.**

DAVID results for gene proliferation phenotypes.

**Table S5.**

Reactome results for gene proliferation phenotypes.

**Table S6.**

Results from proliferation analysis following conserved region disruption.

**Table S7.**

Summary of conserved region disruption followed by amplicon resequencing.

**Table S8.**

JASPAR 2018 TFBS enrichment results within conserved region phenotypes.

**Table S9.**

Enhancer-level summary and proliferation phenotype permutation results.

**Table S10.**

Enhancer-gene chromatin interaction results for proliferation phenotypes.

---

**Fig. S1.**
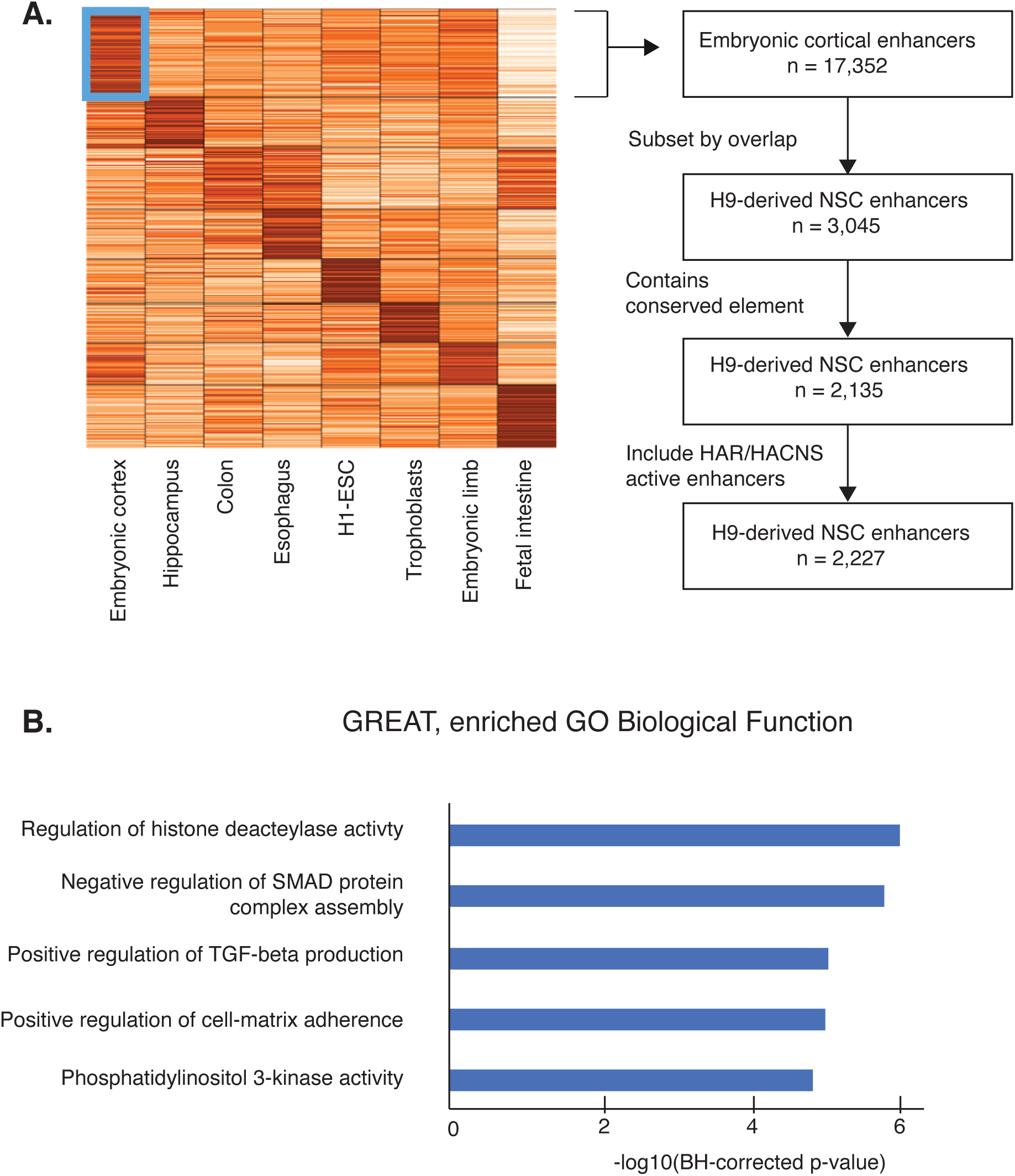
Identification of cortical-enriched enhancers active in hNSC. (**A**) (Left) K-means clustering with embryonic, fetal, and other adult cell and tissue types to identify cortical-associated H3K27ac intervals. (Right) Filtering scheme for the identification of active H3K27ac intervals overlapping active H3K27ac intervals in hNSC. (**B**) Gene ontology enrichments identified by GREAT for cortical-enhancers active in hNSC. P-values are derived from the binomial test reported by GREAT.

**Fig. S2.**
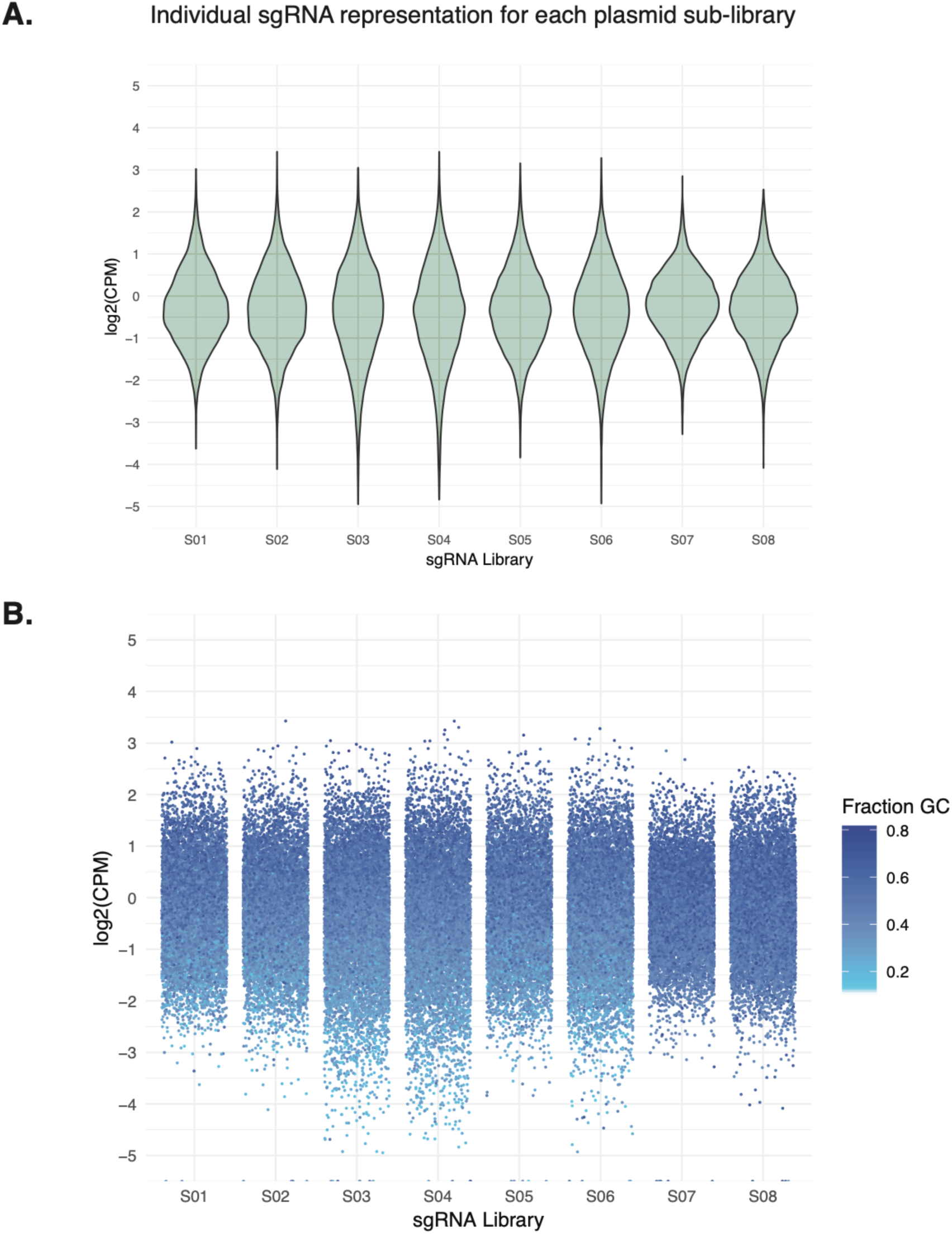
Quality control of sgRNA-Cas9-GFP plasmid libraries. Plasmid libraries S01-S06 are enhancer targeting and S07-S08 are gene. (**A**) Violin plots of individual sgRNA representation relative to mean representation. Log2(counts per million) are shown for each sub-library included in the study. (**B**) Jitter plots showing the percentage GC-content of individual sgRNAs for each sub-library.

**Fig. S3.**
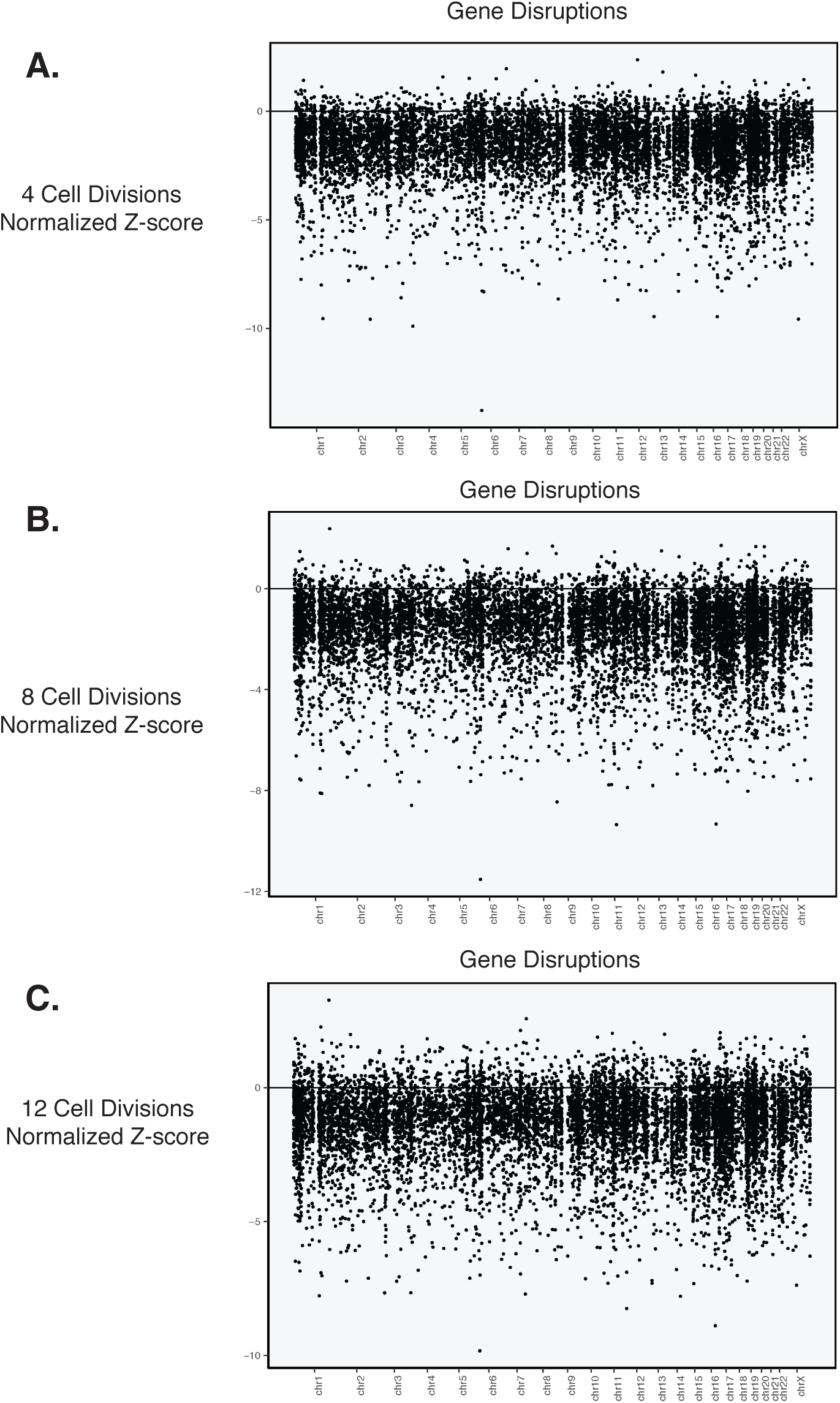
Genome-wide plots of gene disruption biological effects on cell proliferation. The biological effects on cell proliferation are normalized Z-scores relative to mean and standard deviation of the genomic background controls. Plots are shown for all genomic loci at 4 cell divisions (**A**), 8 cell divisions (**B**), and 12 cell divisions (**C**).

**Fig. S4.**
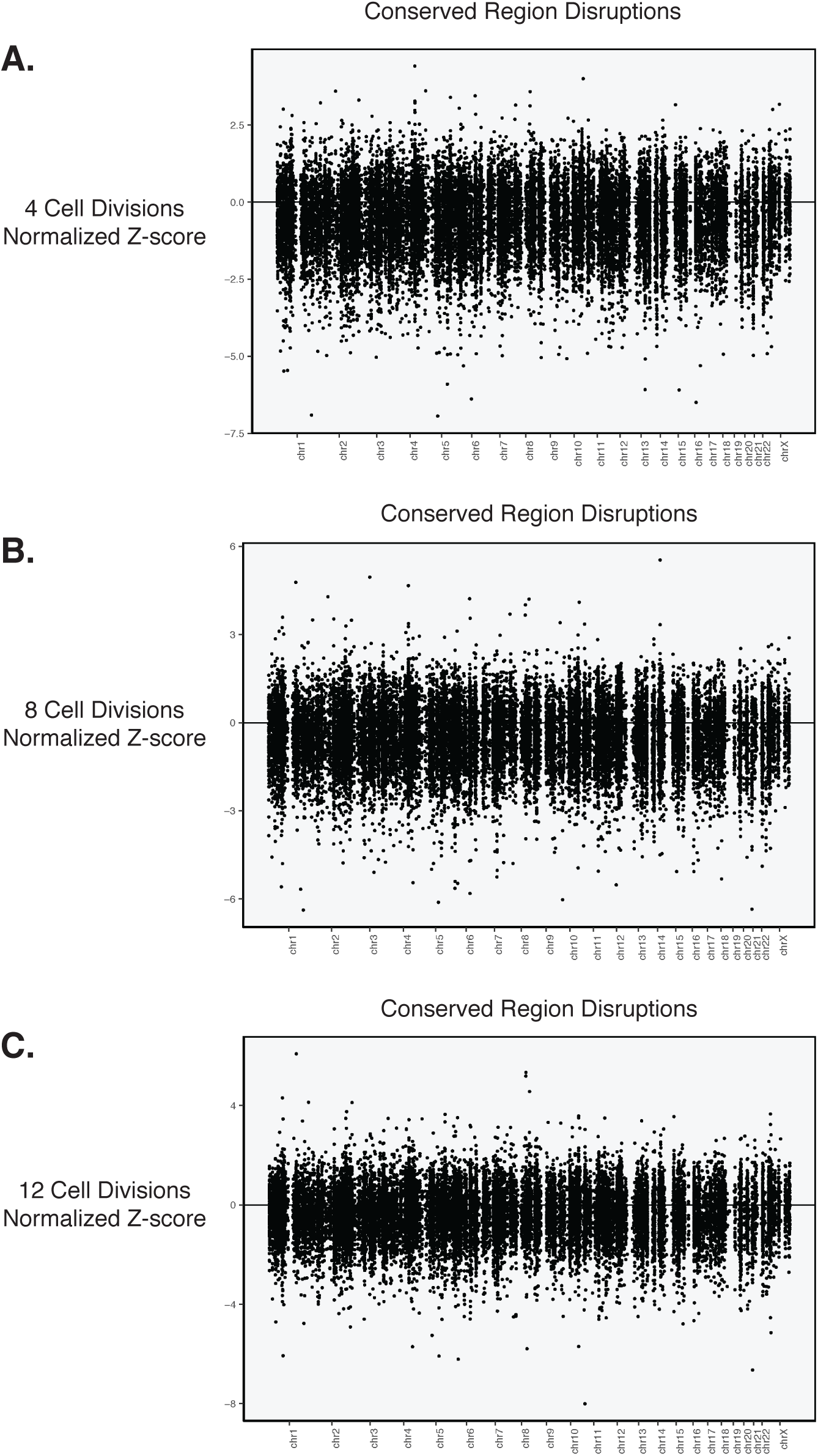
Genome-wide plots of conserved region disruption biological effects on cell proliferation. The biological effects on cell proliferation are normalized Z-scores relative to mean and standard deviation of the genomic background controls. Plots are shown for all genomic loci at 4 cell divisions (**A**), 8 cell divisions (**B**), and 12 cell divisions (**C**).

**Fig. S5.**
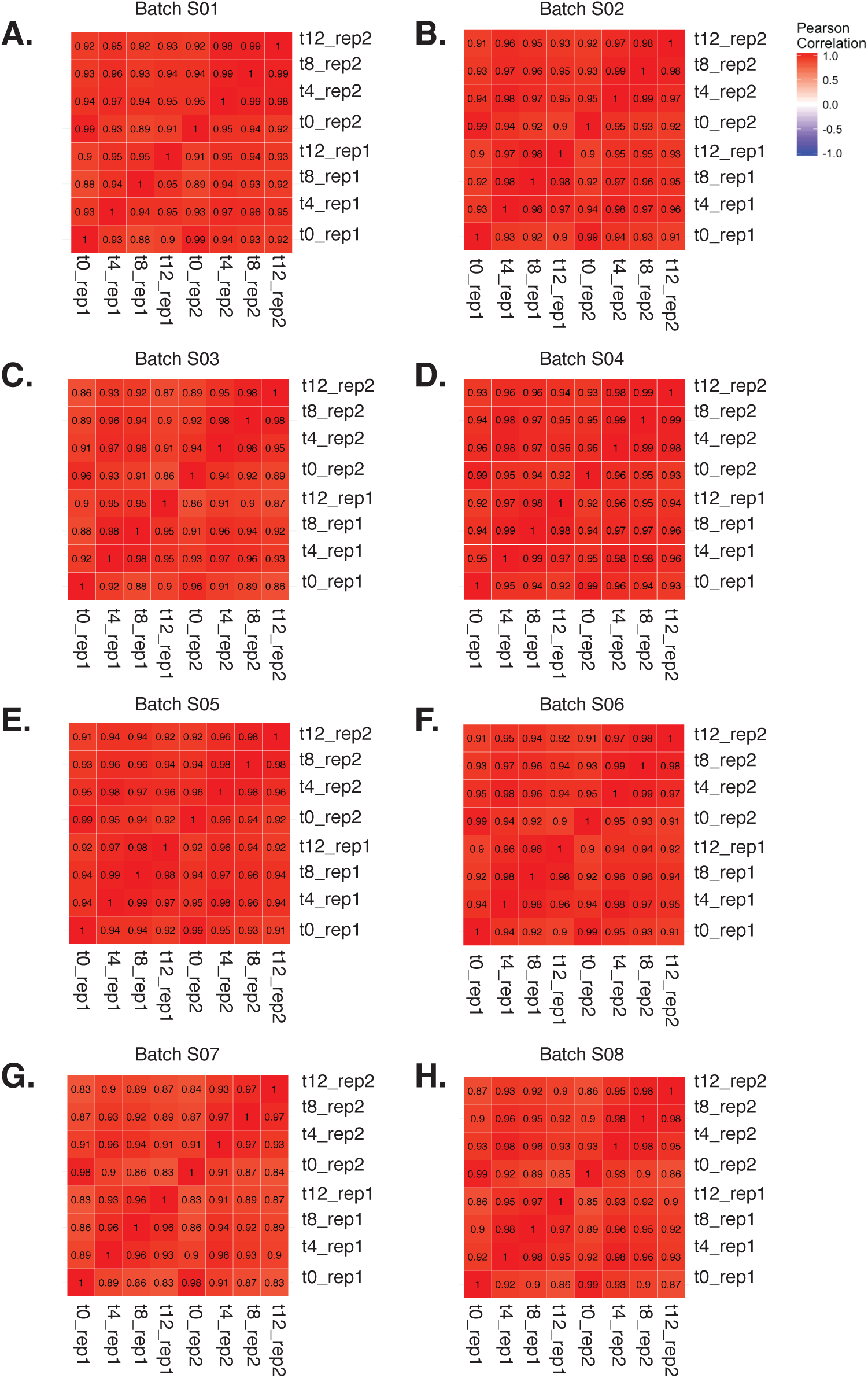
Lentivirus-integrated sgRNA read count correlation plots. (**A-H**) For each sub-library, the Pearson correlation coefficient is reported for t0, t4, t8, and 12 for rep1 and rep2 read counts sequenced and mapped by high-throughput sequencing.

**Fig. S6.**
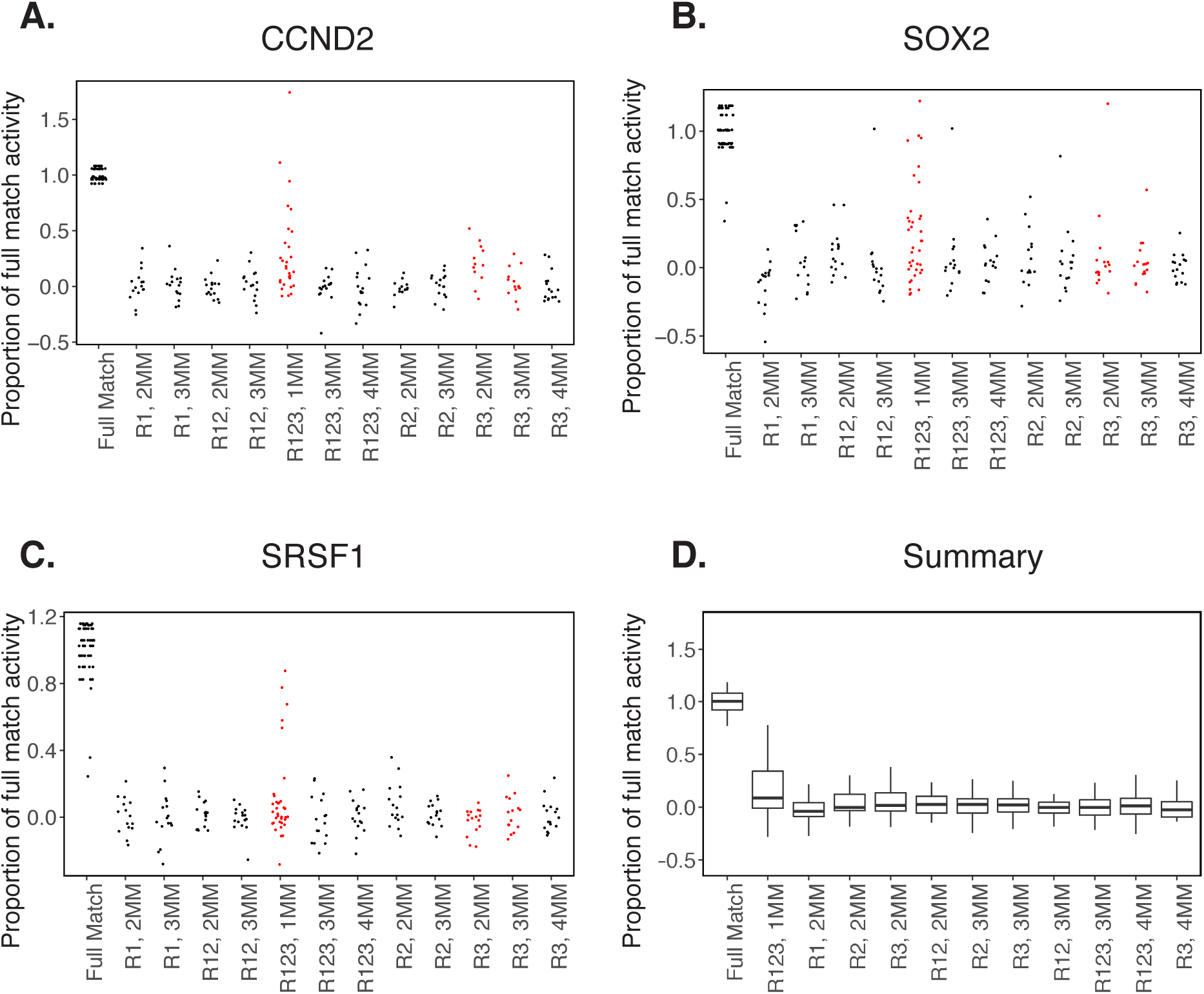
Proportion of on-target activity quantified by specificity controls. The sgRNA scoring scheme was used to test potential off-target effects on cell proliferation. Mismatches were introduced to the on-target sgRNA sequence for CCND2 (**A**), SOX2 (**B**), and SRSF1 (**C**). R1, R2, and R3 are regions of the 20 nt sgRNA sequences where mismatches were introduced. For each condition, 1MM (1 mismatch) 2MM (2 mismatches), 3MM (3 mismatches), or 4MM (4 mismatches) were introduced within the region to generate specificity control sgRNAs (n > 15 sgRNAs per condition). Jitter plots for each region and mismatch pattern are shown (**A-C**). Jitter plots compatible with the scoring criteria to be included in the study (black) and those not meeting scoring criteria (red) are shown. (**D**) Summary of on-target activity across CCND2, SOX2, and SRSF1 when mismatches are introduced.

**Fig. S7.**
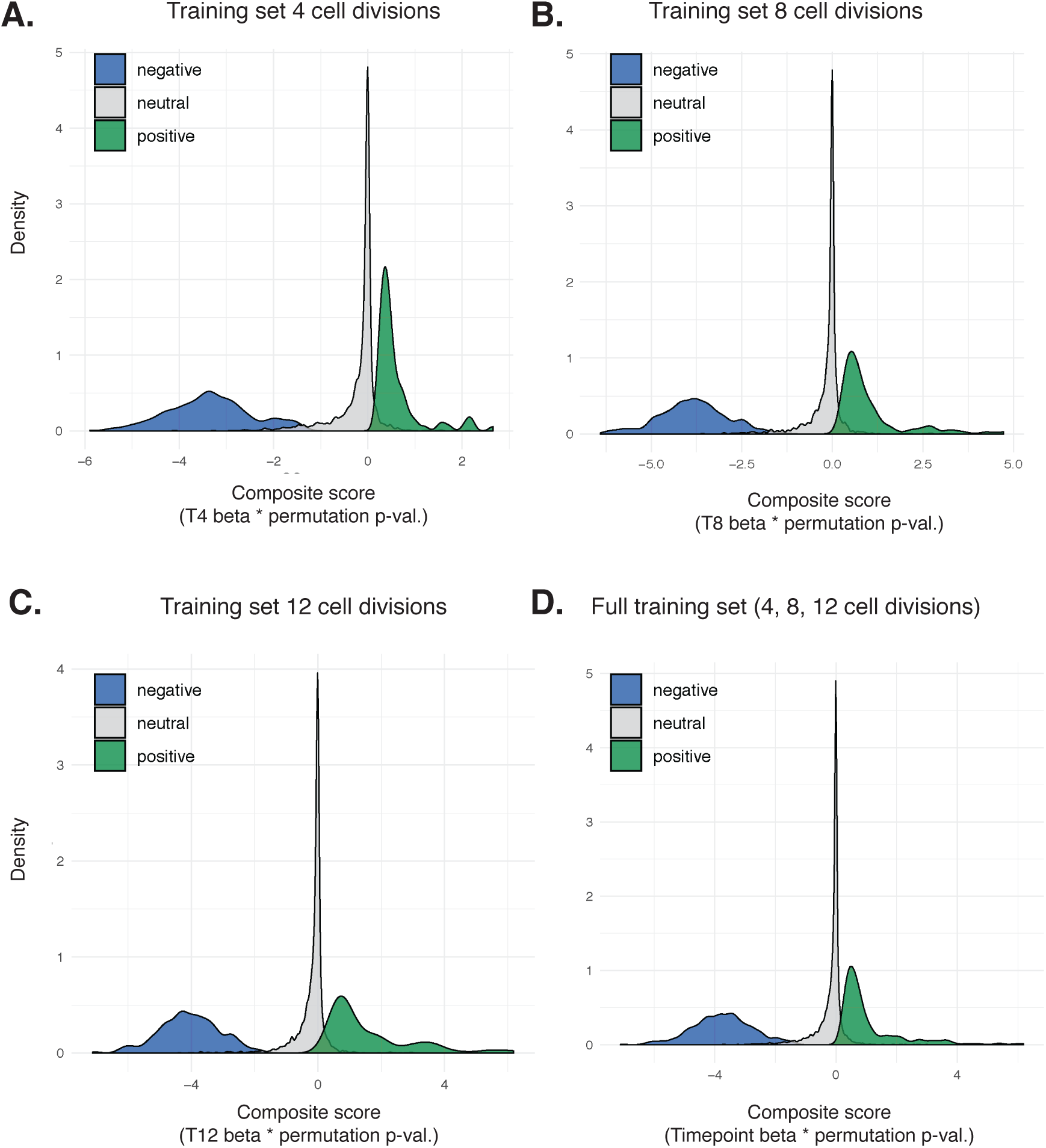
Training set for linear-discriminant analysis and the identification of proliferation phenotypes. Density plots for negative (blue), neutral (gray), and positive (green) proliferation phenotypes are shown for 4 cell divisions (**A**), 8 cell divisions (**B**), and 12 cell divisions (**C**). (**D**) The full training set across all timepoints included in the study.

**Fig. S8.**
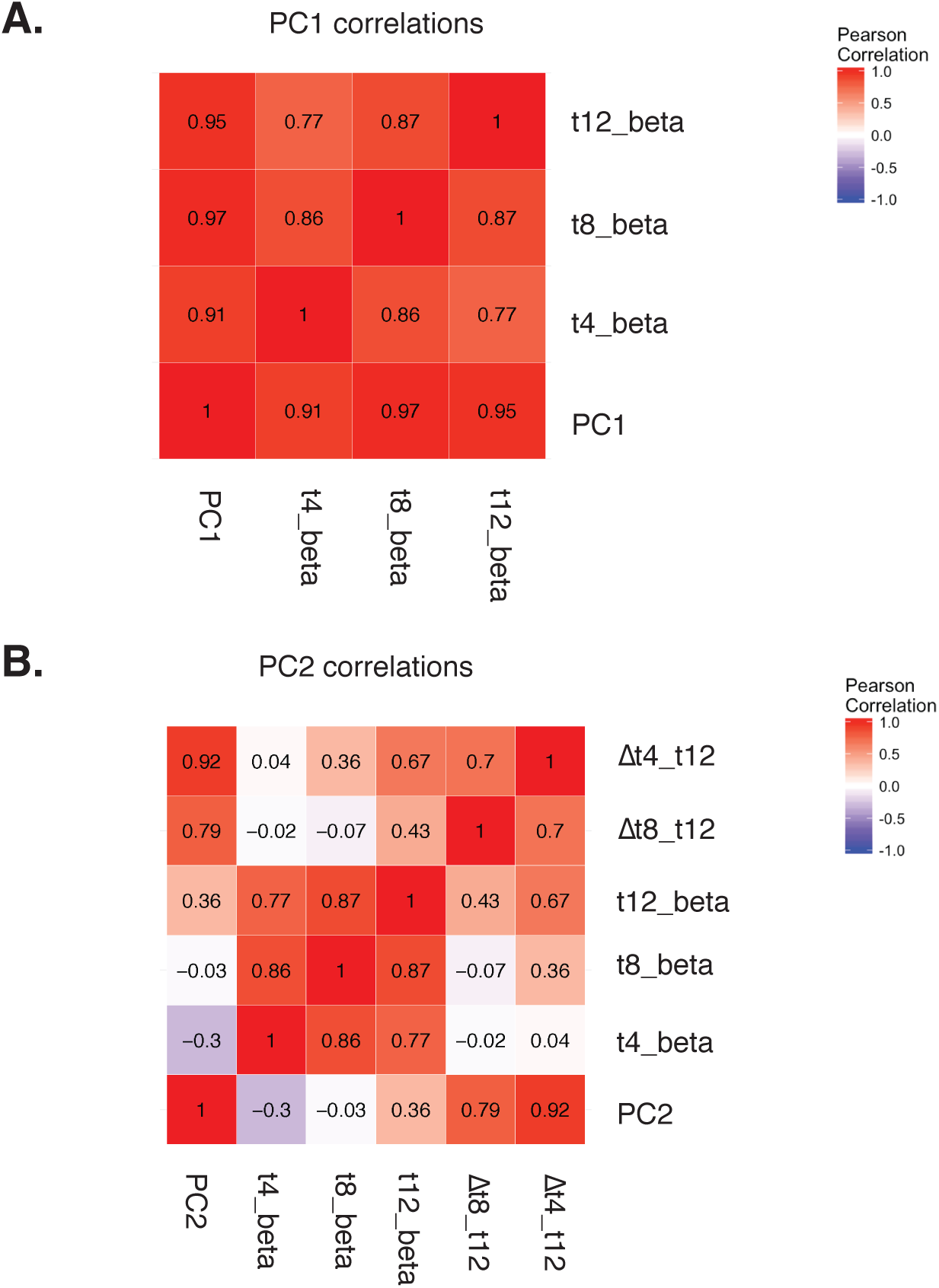
PC1 and PC2 correlation matrices following principal component analysis. (**A**) PC1 Pearson correlations with Beta scores across all targeted regions (genes, enhancers, genomic background controls) and time points. (B) PC2 correlations with joint beta scores and change in beta scores calculated from 8 to 12 cells divisions and from 4 to 12 cell divisions.

**Fig. S9.**
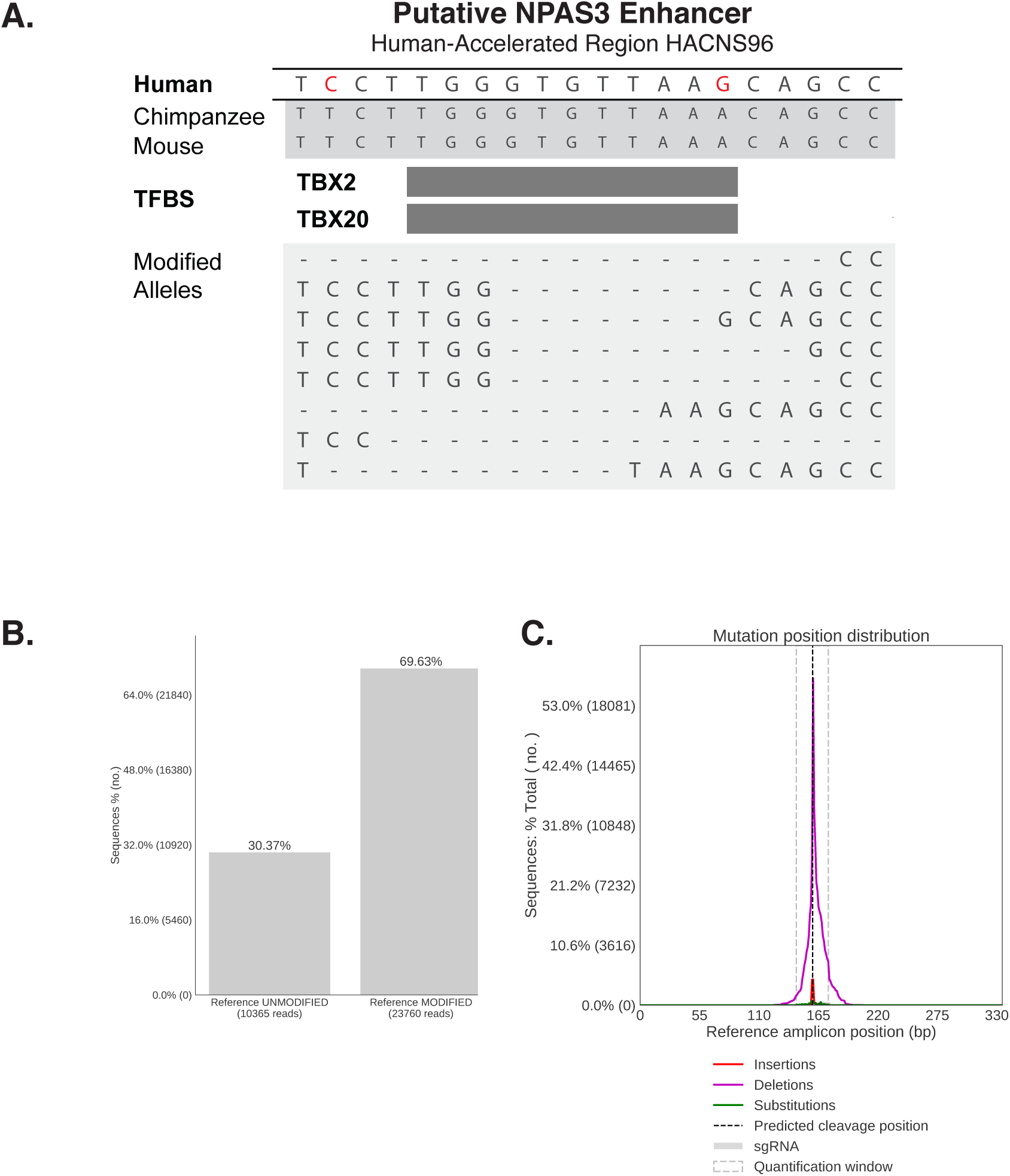
Example of a proliferation-altering disruption that affects human-specific substitutions within a HAR (*HACNS96*). (**A**) (Top) Alignment of *HACNS96* with orthologous sequences in chimpanzee, and mouse, adapted from the 46-way UCSC Multiz alignment (GRCh37/hg19). A human-specific mutation (red) overlaps with predicted TBX2/TBX20 transcription factor binding site (JASPAR 2018). (Bottom) Modified alleles resulting from sgRNA-Cas9-induced disruption. (**B**) Quantification of allele modification following sgRNA-Cas9 disruption. (**C**) Mutational spectrum around the predicted cleavage site for sgRNA-Cas9 disruptions.

**Fig. S10.**
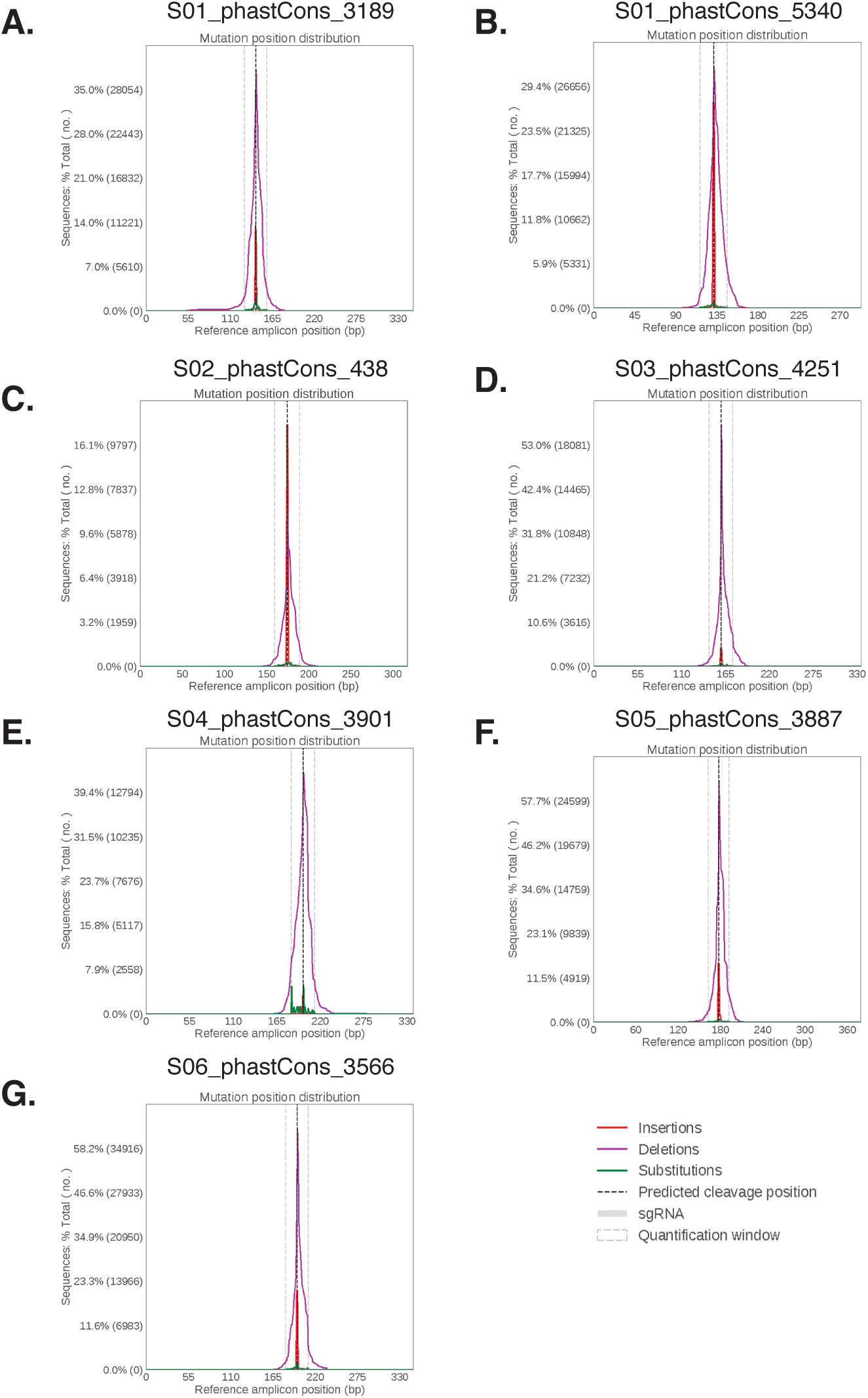
Mutational spectra for disruptions within individual conserved regions. (**A-G**) Mutation types are quantified at the predicted cleavage site for each individual sgRNA targeting the conserved region. Conserved region identifiers and genomic coordinates (GRCh37/hg19) are reported in Table S7.

